# Modeling microRNA-driven post-transcriptional regulation by using exon-intron split analysis (EISA) in pigs

**DOI:** 10.1101/2021.07.14.452370

**Authors:** Emilio Mármol-Sánchez, Susanna Cirera, Laura M. Zingaretti, Mette Juul Jacobsen, Yuliaxis Ramayo-Caldas, Claus B. Jørgensen, Merete Fredholm, Tainã Figueiredo Cardoso, Raquel Quintanilla, Marcel Amills

**Author notes:** **Corresponding author:** Emilio Mármol-Sánchez. Science for Life Laboratory, Department of Molecular Biosciences, The Wenner-Gren Institute. Stockholm University, Stockholm, Sweden. Emilio Mármol-Sánchez current affiliation: ^1^Department of Molecular Biosciences, The Wenner-Gren Institute, Stockholm University, Stockholm, Sweden. ^2^Centre for Paleogenetics, Stockholm University, Stockholm, Sweden. Tainã Figueiredo Cardoso current affiliation: Embrapa Pecuária Sudeste, Empresa Brasileira de Pesquisa Agropecuária (EMBRAPA), 13560-970, São Carlos, SP, Brazil. Emilio Mármol-Sánchez. Claus B. Jørgensen. Tainã Figueiredo Cardoso.

## Abstract

The contribution of microRNAs (miRNAs) to mRNA regulation has often been explored by *post hoc* selection of downregulated genes and determining whether they harbor binding sites for miRNAs of interest. This approach, however, does not discriminate whether these mRNAs are also downregulated at the transcriptional level. Here, we have characterized the transcriptional and post-transcriptional changes of mRNA expression in two porcine tissues: *gluteus medius* muscle of fasted and fed Duroc gilts and adipose tissue of lean and obese Duroc-Göttingen minipigs. Exon-intron split analysis (EISA) of RNA-seq data allowed us to identify downregulated mRNAs with high post-transcriptional signals in fed or obese states, and we assessed whether they harbor binding sites for upregulated miRNAs in any of these two physiological states. We found 26 downregulated mRNAs with high post-transcriptional signals in the muscle of fed gilts and 21 of these were predicted targets of upregulated miRNAs also in the fed state. For adipose tissue, 44 downregulated mRNAs in obese minipigs displayed high post-transcriptional signals, and 25 of these were predicted targets of miRNAs upregulated in the obese state. These results suggest that the contribution of miRNAs to mRNA repression is more prominent in the skeletal muscle system. Finally, we identified several genes that may play relevant roles in the energy homeostasis of the pig skeletal muscle (*DKK2* and *PDK4*) and adipose (*SESN3* and *ESRRG*) tissues. By differentiating transcriptional from post-transcriptional changes in mRNA expression, EISA provides a valuable view about the regulation of gene expression, complementary to canonical differential expression analyses.

## Introduction

The post-transcriptional regulation of gene expression plays a fundamental role towards shaping fine-tuned biological responses to environmental changes (Schaefke *et al*. 2018). Such regulation can take place at multiple levels including splicing, 3′-cleavage and polyadenylation, decay or translation, and its main effectors are RNA binding proteins and non-coding RNAs (Schaefke *et al*. 2018). Of particular importance are microRNAs (miRNAs), which are primarily engaged in the post-transcriptional control of gene expression through inhibition of translation and/or degradation of target mRNAs (Bartel, 2018).

Multiple differential expression studies have been performed in pigs during the last decade (Pérez-Montarelo *et al*. 2013; Óvilo *et al*. 2014; Pilcher *et al*. 2015; Horodyska *et al*. 2018; Benítez *et al*. 2019). One of the main limitations of these studies is that the transcriptional and post-transcriptional components of gene regulation are not independently analyzed. This means that genes that are transcriptionally upregulated and post-transcriptionally downregulated, or vice versa, might not be detected as significantly differentially expressed. Another disadvantage of this approach is that it does not provide insights about the causes of the observed downregulation of RNA transcripts. For instance, studies have typically focused on specific sets of downregulated genes harboring binding sites for miRNAs, in order to disentangle regulatory functions driven by miRNAs (Han *et al*. 2017; Xie *et al*. 2019; Ali *et al*. 2021). This approach, however, cannot distinguish between transcriptional or post-transcriptional repression, which is essential to understand at which level of the mRNA life-cycle regulation is taking place.

To overcome this important limitation, Gaidatzis *et al*. (2015) devised the *exon-intron split analysis* (EISA), which separates the transcriptional and post-transcriptional components of gene regulation by comparing the amounts of exonic and intronic reads from expressed mRNA transcripts. The main assumption of this method is that intronic reads are mainly derived from nascent unprocessed mRNAs or pre-mRNAs, so they reflect transcriptional changes, while post-transcriptional changes can be inferred from differences between the levels of the exonic and intronic fractions (Ameur *et al*. 2011; Gaidatzis *et al*. 2015). For instance, a gene showing little to no differences in the number of sequenced intronic reads, but a strong downregulation of exonic reads after a certain treatment or challenge (nutrition, infection, temperature etc.), could be subjected to post-transcriptional repression (Gaidatzis *et al*. 2015; Cursons *et al*. 2018; Pillman *et al*. 2019). Recent developments on this principle have also been applied to infer the transcriptional fate of cells (La Manno *et al*. 2018).

The main goal of the present study was to investigate the contribution of miRNAs to post-transcriptional regulatory responses using the EISA methodology, combined with *in silico* prediction of miRNA-mRNA interactions and covariation analyses in porcine skeletal muscle and adipose tissues.

## Materials and methods

### Experimental design, sampling and processing

Two experimental systems were used:

(i) Duroc pigs: Twenty-three gilts were subjected to two fasting/feeding regimes, i.e. 11 gilts (*AL-T0*) were slaughtered in fasting condition, while 12 gilts (*AL-T2*) were slaughtered after 7 h of having access to food (Cardoso *et al*. 2017; Ballester *et al*. 2018; Mármol-Sánchez *et al*. 2020). Immediately after slaughtering, *gluteus medius* (GM) skeletal muscle samples were collected and snap-frozen at -80°C.

(ii) Duroc-Göttingen minipig F_2_ inter-cross: Ten individuals fed *ad libitum* with divergent fatness profiles for body mass index (BMI, 5 *lean* and 5 *obese*) were selected from the UNIK resource population (Kogelman *et al*. 2013; Jacobsen *et al*. 2019). Retroperitoneal adipose tissue was collected at slaughter and mature adipocytes were subsequently isolated as previously reported (Jacobsen *et al*. 2019). UNIK minipigs BMI profiles are detailed in **Table S1**.

RNA-seq and small RNA-seq expression data generated in the framework of these two experimental systems have been described previously (Cardoso *et al*. 2017; Jacobsen *et al*. 2019; Mármol-Sánchez *et al*. 2020). Sequencing reads from the RNA-seq and small RNA-seq datasets were trimmed with the Cutadapt software (Martin, 2011). RNA-seq reads were then mapped with the HISAT2 aligner (Kim *et al*. 2019) using default parameters. The Bowtie Alignment v.1.2.1.1 software (Langmead *et al*. 2009) was used to align small RNA-seq reads by considering small sequence reads specifications (*bowtie -n 0 -m 20 -k 1 --best*). The Sscrofa11.1 porcine reference assembly (Warr *et al*. 2020) was used for mapping.

### Exon/Intron quantification

We generated exonic and intronic-specific annotations spanning all available genes by using the Sscrofa.11.1 v.103 gene annotation (Ensembl repositories: http://ftp.ensembl.org/pub/release-103/gtf/sus_scrofa/). Overlapping intronic/exonic regions (10 bp at both ends of introns), as well as singleton positions were removed (Lawrence *et al*. 2013). We then used the *featureCounts* tool (Liao *et al*. 2014) to independently quantify exon and intron expression levels for each mRNA encoding gene, as well as expression levels of mature miRNAs.

### Differential expression analyses

Canonical differential expression analyses were carried out with the *edgeR* tool (Robinson *et al*. 2010) by considering only the exonic counts of mRNAs and the miRNA expression levels measured in the two experimental systems under study. Only genes showing average expression values above 1 count-per-million in at least 50% of animals within each treatment group (*AL-T0* and *AL-T2* for GM tissue and *lean* and *obese* for adipocyte isolates) were retained for further analyses. Expression filtered raw counts for exonic mRNA and miRNA reads were normalized with the trimmed mean of M-values normalization (TMM) method (Robinson & Oshlack 2010) and the statistical significance of mean expression differences was tested with a quasi-likelihood F-test (Robinson *et al*. 2010). Multiple hypothesis testing correction was implemented with the false discovery rate method (Benjamini & Hochberg 1995). Messenger RNAs were considered to be significantly differentially expressed when the absolute value of the fold-change (FC) was higher than 2 (|FC| > 2) and the *q*-value < 0.05. Fasting Duroc gilts (*AL-T0*) as well as UNIK *obese* minipigs were classified as baseline controls in differential expression analyses, i.e. any given upregulation in exonic counts represents and overexpression of the corresponding gene in fed (*AL-T2*) Duroc gilts or *lean* UNIK minipigs with respect to their fasting (*AL-T0*) and *obese* counterparts, respectively.

Since changes in the expression of miRNAs are often subtler than those of mRNAs, the thresholds to consider a miRNA as truly differentially expressed were set to |FC| > 1.5 and *q*-value < 0.05 (Guo *et al*. 2015) for fasted (*AL-T0*) vs fed (*AL-T2*) Duroc gilts. Given the relatively low statistical significance of miRNA expression changes observed in *obese* vs *lean* minipigs, we considered, as potential miRNA regulators, those that were significantly (FC < -1.5; *q*-value < 0.05) and suggestively (FC > 1.5 and *P*-value < 0.01) upregulated in *lean* minipigs.

### Exon/intron split analysis (EISA)

We applied EISA to differentiate transcriptional from post-transcriptional changes in mRNA expression in our two experimental systems (muscle and fat). Normalization was performed independently for exon and intron counts as described by Gaidatzis *et al*. (2015). Exonic and intronic gene abundances were subsequently log_2_ transformed, adding a pseudo-count of 1 and averaged within each considered treatment group.

Only genes for which both exonic and intronic read counts were successfully quantified were further considered. Observed differences in each *i*^*th*^ gene were expressed as the increment of exonic/intronic counts in fed (*AL-T2*) and *obese* animals with respect to fasting (*AL-T0*) and *lean* animals, respectively. In this way, the increment of intronic and exonic counts was calculated considering ΔInt = Int_2i_ – Int_1i_ and ΔEx = Ex_2i_ – Ex_1i_, respectively. The magnitudes of the transcriptional (Tc) and post-transcriptional (PTc) changes in mRNA expression were then calculated. The Tc contribution to the observed counts is explained by ΔInt (Pillman *et al*. 2019), while PTc can be deduced from ΔEx – ΔInt. In this way, the significance of Tc scores was assessed as in canonical differential expression analyses but using the intronic fraction as input. Besides, for assessing the statistical significance of PTc scores, we incorporated the fraction type (exonic or intronic counts) as an interaction term with the condition type (fasted *AL-T0* vs fed *AL-T2* or *obese* vs *lean*) in the framework of the generalized linear model (Gaidatzis *et al*. 2015) and using *edgeR* tool with a quasi-likelihood F-test as in differential expression analyses (Robinson *et al*. 2010). Both Tc and PTc components were z-scored to make ΔEx and ΔInt estimates comparable. Moreover, Tc and PTc scores were considered significant when |FC| > 2 and *q*-value < 0.05. Multiple hypothesis testing correction was implemented by using the false discovery rate approach (Benjamini & Hochberg 1995).

In order to obtain a prioritized list of genes showing high signals of post-transcriptional regulation, the top 5% of expressed genes with the most negative PTc scores were retrieved (irrespective of their statistical significance after multiple testing correction). From these, we only focused on genes showing strongly reduced ΔEx values of at least 2-fold for post-transcriptional signals in both experimental systems (i.e. ΔEx < -1 in the log_2_ scale). All implemented analyses have been summarized in **Fig. S1**. A ready-to-use modular pipeline for running EISA is publicly available at https://github.com/emarmolsanchez/EISAcompR.

### miRNA target prediction

Putative interactions between the seed of expressed miRNAs (2^nd^-8^th^ 5’ nts) and the 3’-UTRs of expressed mRNA genes were predicted on the basis of sequence identity using the Sscrofa11.1 reference assembly and the seedVicious v1.1 tool (Marco 2018). The annotated 3’-UTRs longer than 30 nts from porcine mRNAs were retrieved from http://www.ensembl.org/biomart, while mature miRNA sequences were obtained from miRBase (Kozomara *et al*. 2019). Redundant seeds from mature porcine microRNAs were removed, and 8mer, 7mer-m8 and 7mer-A1 miRNA-mRNA interactions were taken into account (Bartel 2018).

Based on Grimson *et al*. (2007), *in silico*-predicted miRNA-mRNA interactions matching any of the following criteria were removed: (i) Binding sites located in 3’-UTRs at less than 15 nts close to the end of the open reading frame (and the stop codon) or less than 15 nts close to the terminal poly(A) tail (E criterion), (ii) binding sites located in the middle of the 3’-UTR in a range comprising 45-55% of the central region of the non-coding sequence (M criterion), and (iii) binding sites that lack AU-rich sequences in their immediate upstream and downstream flanking regions comprising 30 nts each (AU criterion). A schematic representation of such criteria is available at **Fig. S2**.

Covariation patterns between miRNAs and their predicted mRNA targets were assessed by computing Spearman’s correlation coefficients (*ρ*) with the TMM normalized and log_2_ transformed expression profiles of the exonic fraction of mRNA and miRNA genes. Only miRNA-mRNA predicted pairs comprising significantly upregulated miRNAs and downregulated mRNA genes with high PTc scores (see exon/intron split analysis section) were considered.

### miRNA target enrichment analyses

We predicted which downregulated mRNA genes, from those with highly negative post-transcriptional signals, are putatively targeted by at least one significantly upregulated miRNA. Subsequently, we investigated whether the sets of mRNA genes identified in this way are more enriched in binding sites for upregulated miRNAs than the whole set of expressed mRNAs genes with available 3’-UTRs (control background). Enrichment analyses were carried out using the Fisher’s exact test in R. Significance level was set at a nominal *P*-value < 0.05.

We also tested whether downregulated mRNA genes with highly negative post-transcriptional signals were significantly enriched to be targets of at least one of the top 5% most highly expressed miRNA genes (considering their overall average expression and excluding significantly upregulated miRNAs), as well as of significantly downregulated miRNAs.

As an additional control randomized test for enrichment analyses between miRNAs and downregulated mRNAs with highly negative post-transcriptional signals, we implemented a bootstrap corrected iteration to generate 100 random sets of 10 expressed mature miRNA genes without seed redundancy. In this way, we predicted which downregulated mRNA genes, from those with highly negative post-transcriptional signals, are putatively targeted by at least one miRNA of the random sets generated. The distribution of odds ratios obtained after enrichment tests over each random set of miRNAs (N = 100) were then compared with the observed odds ratios obtained with the set of significantly upregulated miRNAs.

The *P*-value for the significance of the deviation of observed odds ratios against the bootstrapped odds ratios distribution was defined as:

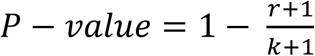, where *r* is the number of permuted odds ratios with values equal or higher than the observed odds ratio, and *k* is the number of permutations (N = 100).

### Gene covariation network and covariation enrichment score

We used *edgeR* to identify differentially expressed genes in the *AL-T0* vs *AL-T2* (N = 454 genes) and *obese* vs *lean* (N = 299 genes) comparisons showing *q*-value < 0.05 after multiple testing correction. Then, the normalized exonic and intronic estimates in the log_2_ scale obtained from EISA analyses were independently used to compute Spearman’s correlation coefficients (*ρ*). Significant correlations were identified with the Partial Correlation with Information Theory (PCIT) algorithm (Reverter & Chan 2008) implemented in the *pcit* R package (Watson-Haigh *et al*. 2010). In this way, the number of overall significant pairwise correlations was compared with that obtained when only considering the set of downregulated mRNA genes with highly negative post-transcriptional signals and putatively targeted by significantly upregulated miRNAs. Thus, the potential contribution of miRNAs to the observed differences in covariation among the two sets of genes was assessed by calculating a covariation enrichment score (CES), as reported by Tarbier *et al*. (2020). Further details about the algorithm used to calculate the CES values and control tests can be found in **Supplementary Methods**. Significant differences among the set of exonic, intronic and control CES values were tested with a Mann-Whitney U non-parametric test (Mann & Whitney 1947).

### Verification of RNA-seq expression profiles by qPCR

By using qPCR, we have verified that the expression profiles of selected mRNAs and miRNAs highlighted in the adipose tissue experiment were in accordance with those obtained with RNA-seq and small RNA-seq data. Since this experiment is just a control of the quality of expression estimates, further details are reported in **Supplementary Methods**. Primers for mRNA and miRNA qPCR expression profiling are available at **Table S2**. Raw Cq values for each assay are available at **Table S3**.

## Results

### The analysis of post-transcriptional regulation in muscle samples from fasting and fed Duroc gilts

#### Differential expression and EISA analyses

Total RNA and small RNA were independently sequenced in muscle samples from fasting (*AL-T0*) and fed (*AL-T2*) Duroc gilts. About 45.2 million reads (93%) per sample from protein coding and non-coding genes were successfully mapped against the Sscrofa.11.1 assembly when analyzing the GM muscle total RNA-seq data. Besides, around 2.2 million reads per sample (42%) obtained from the small RNA-seq experiment were successfully mapped to 370 annotated porcine miRNA genes.

A total of 30,322 (based on exonic reads) and 22,769 (based on intronic reads) genes were successfully quantified after splitting the reference genome assembly between exonic and intronic features. Exonic counts were ∼23 fold more abundant than those corresponding to intronic regions.

By using *edgeR*, we detected 454 mRNA genes with *q*-value < 0.05 (**Table S4a**). Among these, only genes with |FC| > 2 were retained, resulting in 52 upregulated and 80 downregulated genes (**Table S4a** in bold and **Fig. S3a**). The analysis of small RNA-seq data with *edgeR* revealed 16 miRNAs significantly differentially expressed, of which 8 were upregulated in *AL-T2* gilts. These 8 miRNAs, which represented 6 unique miRNA seeds (ssc-miR-148a-3p, ssc-miR-7-5p, ssc-miR-30-3p, ssc-miR-151-3p, ssc-miR-374a-3p and ssc-miR-421-5p; **Table S5** in bold), were selected as potential post-transcriptional regulators of mRNA genes.

On the other hand, EISA highlighted 26 mRNA genes displaying the top 5% negative PTc scores with at least 2-fold ΔEx reduction (**Table 1** and **Fig. S3b**). Eighteen out of these 26 genes (69.23%) appeared as significantly downregulated (FC < -2; *q*-value < 0.05, **Table 1** and **Table S4b** in bold) according to differential expression analyses. Also, we detected 133 mRNA genes with significant PTc scores (|FC| > 2; *q*-value < 0.05, **Table S6a**), of which three experienced at least a 2-fold reduction of their ΔEx fraction (**Table S6a** in bold). Two out from these three mRNA genes ranked within those with the top 5% negative PTc scores (**Table 1** and **Table S6a**). Among this set of 133 genes, only seven (5.26%) were also significantly differentially expressed (**Table S6a**).

**Table 1:**
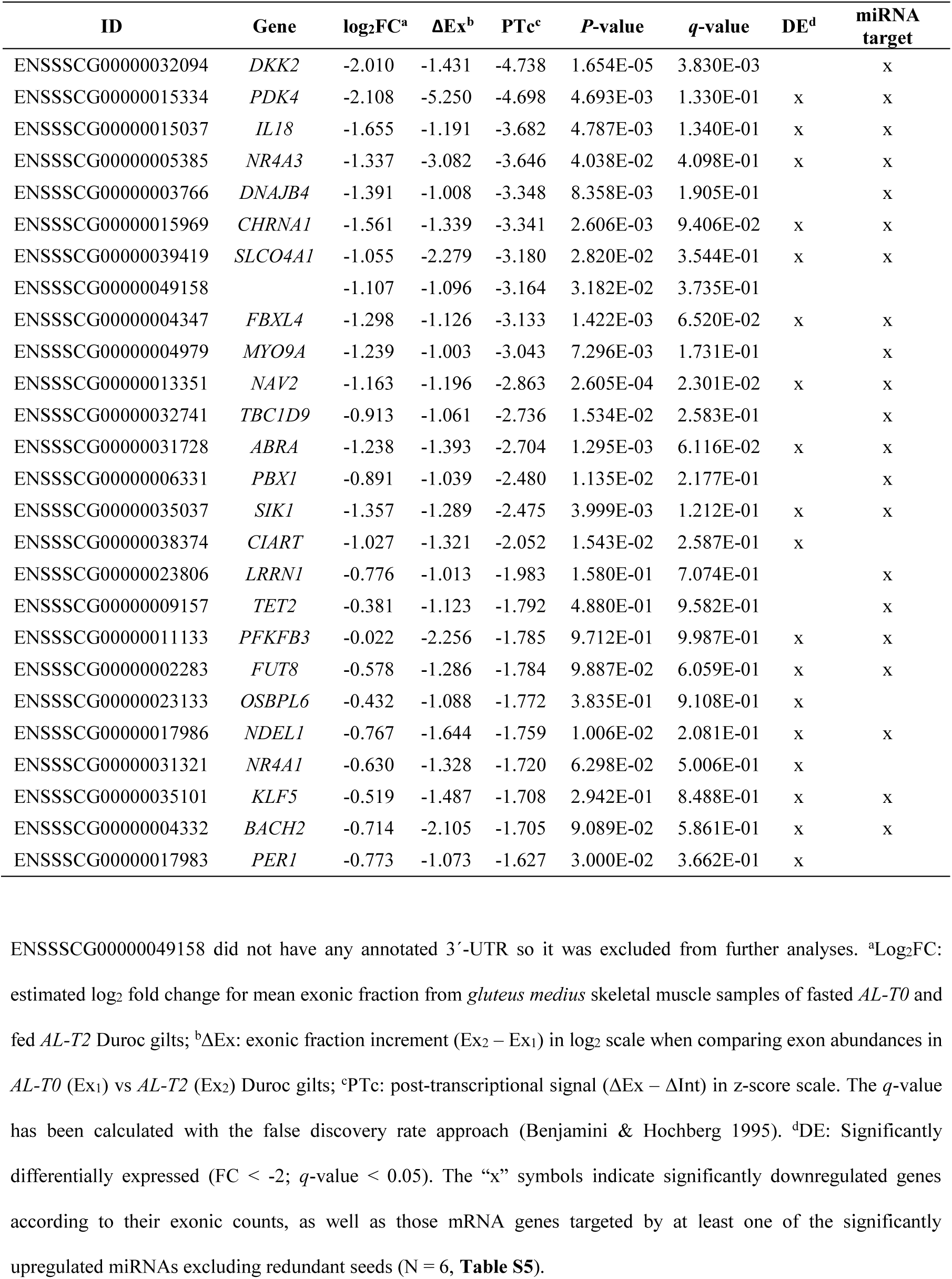
mRNA genes with the top 5% post-transcriptional (PTc) scores and at least 2-fold exonic fraction (ΔEx) reduction (equivalent to -1 in the log_2_ scale) of *gluteus medius* skeletal muscle samples from fasting (*AL-T0*, N = 11) and fed (*AL-T2*, N = 12) Duroc gilts.

Moreover, with EISA we detected 344 genes displaying significant Tc scores (|FC| > 2; *q*-value < 0.05, **Table S6b**) and among these, 71 (20.63%) were also significantly differentially expressed (**Tables S6b**). Besides, 91 out of these 344 genes (26.45%) also showed significant PTc scores (**Table S6b** in bold), but none of them were among the mRNA genes displaying the top 5% negative PTc scores with at least 2-fold ΔEx reduction (N = 26, **Table 1**). The whole list of expressed mRNA genes after differential expression analyses is available at **Table S4c**. The whole lists of expressed mRNA genes after EISA and their PTc and Tc scores are available at **Table S6c** and **S6d**, respectively.

#### Context-based pruning of predicted miRNA-mRNA interactions removes spurious unreliable target events

Before making *in silico* predictions of miRNA-mRNA interactions, we investigated their reliability (**Table S7**). To do so, we first evaluated the enrichment in the number of genes with binding sites for at least one of the 6 non-redundant miRNAs upregulated in the GM muscle of *AL-T2* gilts (ssc-miR-148a-3p, ssc-miR-7-5p, ssc-miR-30-3p, ssc-miR-151-3p, ssc-miR-374a-3p and ssc-miR-421-5p) over a random background of expressed genes with no context-based removal of predicted binding sites (see Methods). Introducing additional context-based filtering criteria to remove unreliable binding site predictions resulted in an overall increased enrichment of target genes within the list of the top 1% (N = 13 genes, **Fig. S4a**) and 5% (N = 26 genes, **Fig. S4b**) genes with negative PTc scores and displaying at least 2-fold ΔEx reduction. This significant enrichment was more evident when using the AU criterion, as shown in **Fig. S4a**.

#### Several genes with relevant post-transcriptional signals detected with EISA are predicted to be targets of upregulated miRNAs

Target prediction and context-based pruning of miRNA-mRNA interactions for mRNA genes displaying the top 5% negative PTc scores and at least 2-fold reduction in the ΔEx exonic fraction (N = 26, **Table 1, Fig. 1a**) made possible to detect 11 8mer, 21 7mer-m8 and 22 7mer-A1 miRNA binding sites for the six non-redundant seeds of miRNAs significantly upregulated in *AL-T2* gilts (**Table S5** in bold) in 21 out of the 26 analyzed mRNAs (80.77%, **Table S7**). Moreover, 14 out of these 21 genes (66.67%) were also significantly differentially expressed (**Table 1** and **Table S4b** in bold).

**Figure 1:**
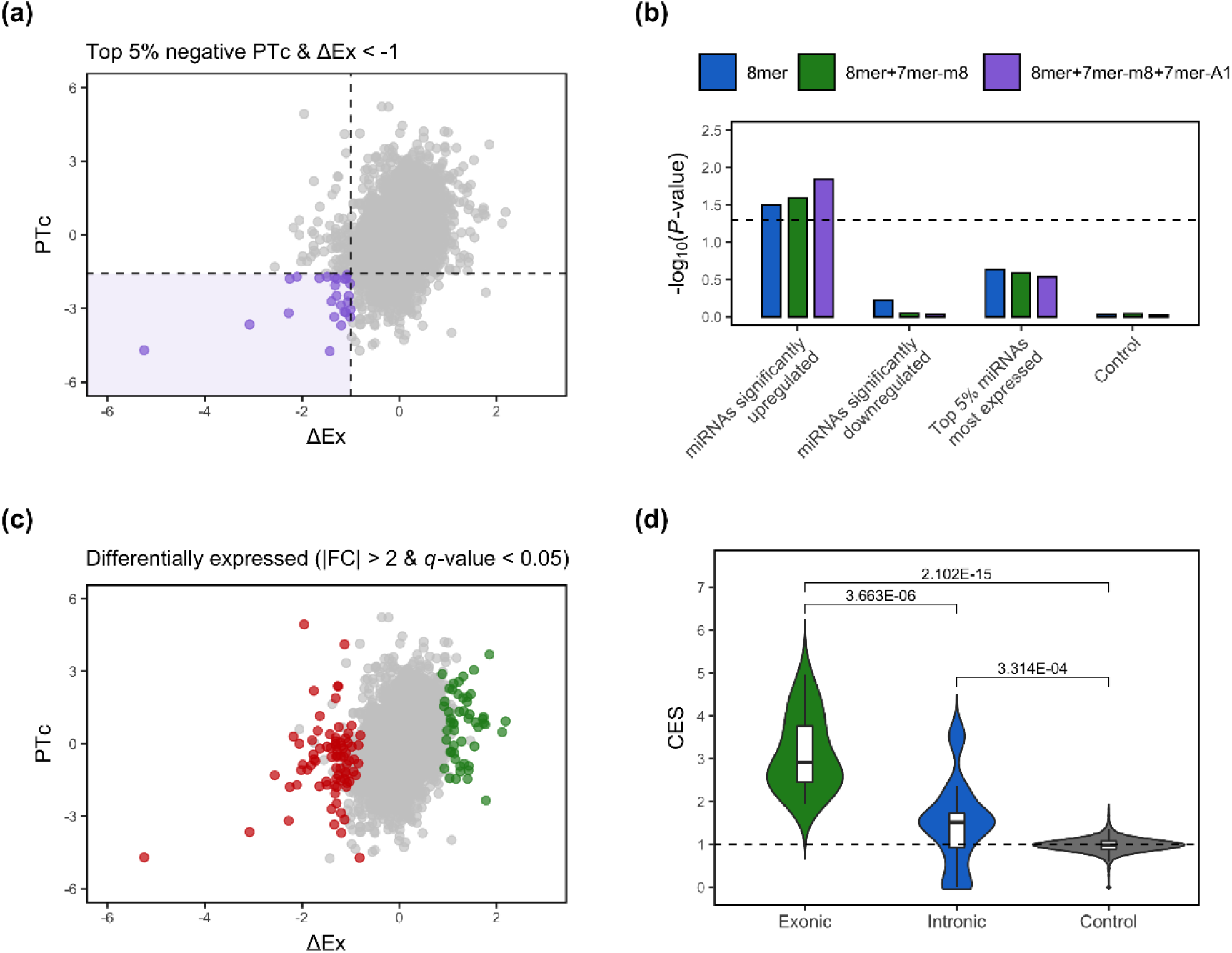
(**a**) Scatterplot depicting genes expressed in *gluteus medius* skeletal muscle of fasted (*AL-T0*, N = 11) and fed (*AL-T2*, N = 12) Duroc gilts according to their exonic fraction (ΔEx) and post-transcriptional (PTc) scores. Genes with the top 5% negative PTc scores and at least 2-fold ΔEx reduction (equivalent to -1 in the log_2_ scale) are highlighted in purple and delimited by dashed lines. (**b**) Enrichment analyses comparing all expressed mRNA genes and the set of mRNA genes with the top 5% negative PTc scores and at least 2-fold ΔEx reduction as being putatively targeted by either significantly upregulated miRNAs (FC > 1.5; *q*-value < 0.05), significantly downregulated miRNAs (FC < -1.5; *q*-value < 0.05) or the top 5% most highly expressed miRNAs, excluding significantly upregulated miRNAs. As indicated with the dashed line, a nominal *P*-value = 0.05 was set as a significance threshold. (**c**) Scatterplot depicting genes expressed in *gluteus medius* skeletal muscle of fasted (*AL-T0*, N = 11) and fed (*AL-T2*, N = 12) Duroc gilts according to their exonic fraction (ΔEx) and post-transcriptional (PTc) scores. Genes significantly upregulated are in green, while those being downregulated are in red (|FC| > 2; *q*-value < 0.05). (**d**) Covariation enrichment scores (CES) for the exonic and intronic fractions of the 21 mRNA genes with the top 5% negative PTc scores and at least 2-fold ΔEx reduction predicted to harbor binding sites for upregulated miRNAs (N = 6) in the *gluteus medius* skeletal muscle of fasted (*AL-T0*, N = 11) and fed (*AL-T2*, N = 12) Duroc gilts. The control set was established by generating 1,000 permuted lists of 21 genes chosen at random and using their exonic and intronic fractions for calculating their CES values. Statistical significance was assessed using a Mann-Whitney U non-parametric test (Mann & Whitney 1947). The dashed line represents a CES of 1, equivalent to an observed null fold change in covariation.

This set of 21 mRNA genes with putative post-transcriptional repression mediated by miRNAs showed a significant enrichment in 8mer, 7mer-m8 and 7mer-A1 sites for the 6 miRNAs significantly upregulated in *AL-T2* gilts, especially when combining these three types of binding sites altogether (**Fig. 1b**). The miRNAs with the highest number of significant miRNA-mRNA interactions were ssc-miR-30a-3p and ssc-miR-421-5p, which showed nine and eight significant interactions, followed by ssc-miR-148-3p with four significant interactions (**Table S7**).

We also evaluated the enrichment of the mRNA genes within the list of the top 5% negative PTc scores and at least 2-fold ΔEx reduction (N = 26, **Table 1**) to be targets of at least one of the following: (i) miRNAs downregulated in *AL-T2* fed gilts (**Table S5**), (ii) top 5% most expressed miRNAs, excluding those significantly upregulated (ssc-miR-1, ssc-miR-133a-3p, ssc-miR-26a, ssc-miR-10b, ssc-miR-378, ssc-miR-99a-5p, ssc-miR-27b-3p, ssc-miR-30d, ssc-miR-486 and ssc-let-7f-5p), and (iii) random sets (N = 100) of 10 expressed miRNAs, as a control test. In none of these three analyses a significant enrichment was detected (**Fig. 1b**).

The mRNA with the highest negative and most significant PTc score was the Dickkopf WNT Signaling Pathway Inhibitor 2 (*DKK2*), being a strong candidate to be repressed by miRNAs (**Table 1**). Indeed, *DKK2* was the only gene harboring two miRNA 8mer binding sites (**Table S7**). Interestingly, this locus was almost significantly differentially expressed (**Table 1** and **Table S4b**). The discordance between EISA and differential expression results can be fully appreciated when comparing **Fig. 1a** (genes with high post-transcriptional repression after EISA) and **Fig. 1c** (differential expression analyses), where only 18 out of the 26 mRNA genes detected with EISA appeared as significantly downregulated in the *edgeR*-based differential expression analyses (**Table 1**). Although several of the mRNA genes shown in **Table 1** were highly downregulated (**Table S4b**), the majority were mildly to slightly downregulated or not significantly differentially expressed.

#### Genes showing post-transcriptional regulatory signals predominantly covary at the exonic level

To further elucidate whether the set of 21 mRNA genes with putative post-transcriptional repression mediated by upregulated miRNAs showed covarying expression profiles, we evaluated the number of significant co-expressed pairs among them and among the whole set of mRNA genes with *q*-value < 0.05, including those from the set of 21 mRNA genes with *q*-value > 0.05 after differential expression analyses (**Table 4b**).

By calculating CES values for both exonic and intronic fractions (see Methods) of the 21 genes putatively targeted by the 6 significantly upregulated miRNAs, our analyses revealed that 19 out of these 21 genes showed increased covariation in their exonic fraction when compared to their intronic fraction (**Table S8, Fig. 1d**), and *DKK2* was again the gene with the strongest exonic covariation fold change compared to its intronic covariation (**Table S8**). As expected, control random sets of genes (N = 1,000) displayed CES ≈ 1, indicative of no covariation (**Fig. 1d**). The observed CES distributions of exonic and intronic sets were significantly different (*P*-value = 3.663E-06) after running non-parametric tests (**Fig. 1d**), thus supporting that the majority of these 21 genes might be indeed co-regulated at the post-transcriptional level by upregulated miRNAs.

### Studying post-transcriptional signals in adipose tissue using the UNIK minipig population

After pre-processing and filtering of sequenced reads from adipocyte samples, we were able to retrieve ∼98.1 and ∼0.87 million mRNA and small RNA reads per sample, and ∼96.5% and ∼73.4% of these reads mapped to annotated porcine mRNA and miRNA genes, respectively. Differential expression analyses revealed a total of 299 genes with *q*-value < 0.05, of which 52 and 95 were significantly downregulated and upregulated (FC > |2|; *q*-value < 0.05), respectively (**Table S9a** in bold). Only one miRNA (ssc-miR-92b-3p) was significantly upregulated in *lean* minipigs (**Table S10**), while six additional miRNAs showed suggestive differential expression (*P*-value < 0.01), of which four were upregulated (ssc-miR-148a-3p, ssc-miR-204, ssc-miR-92a and ssc-miR-214-3p; **Table S10** in bold).

After running EISA, only the sestrin 3 (*SESN3*) gene showed a significant PTc score, having the second highest negative PTc score (**Table 2** and **Table S11a**). Moreover, *SESN3* was also detected as the most significantly downregulated gene by *edgeR* (**Table S9a**). A total of 44 downregulated mRNAs in *lean* minipigs displayed the top 5% PTc scores with reduced ΔEx of at least 2-fold (**Table 2** and **Fig. 2a**). Among them, only 12 (27.27%) appeared as significantly downregulated (FC < -2; *q*-value < 0.05, **Table 2** and **Table S9b** in bold). The whole list of expressed genes after differential expression analyses are available at **Table S9c**.

**Table 2:**
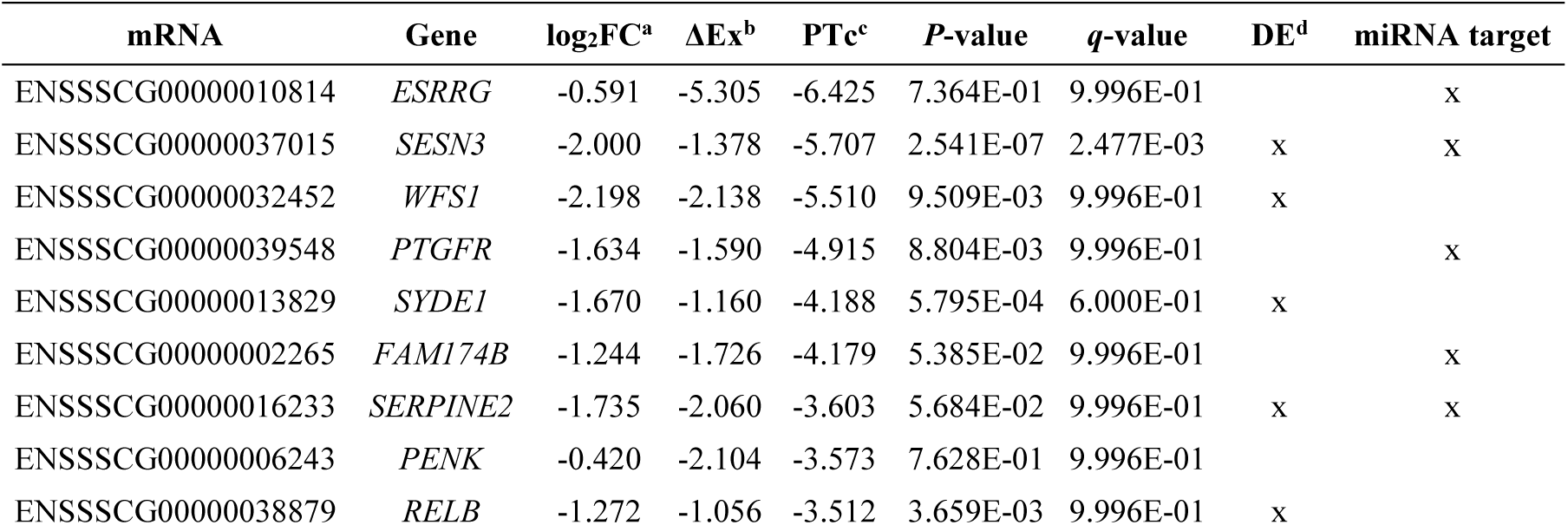
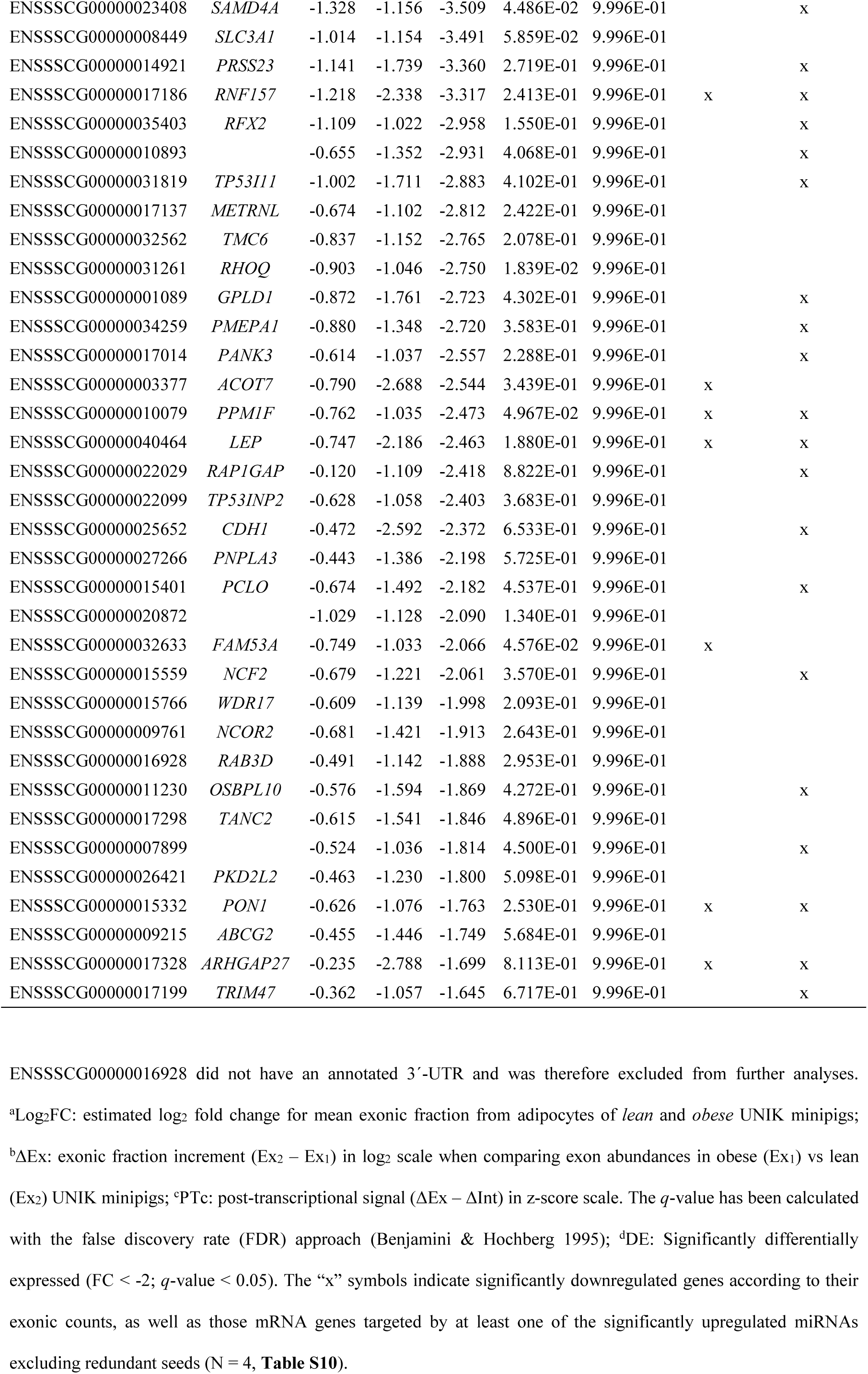
mRNA genes with the top 5% post-transcriptional (PTc) scores and at least 2-fold exonic fraction (ΔEx) reduction (equivalent to -1 in the log2 scale) of adipocytes from *lean* (N = 5) and *obese* (N = 5) UNIK minipigs classified in accordance with their body mass index.

| mRNA | Gene | log<sub>2</sub>FC<sup>a</sup> | $\Delta Ex^b$ | PTc<sup>c</sup> | P-value | q-value | DE<sup>d</sup> | miRNA target |
| --- | --- | --- | --- | --- | --- | --- | --- | --- |
| ENSSSCG00000010814 | *ESRRG* | -0.591 | -5.305 | -6.425 | 7.364E-01 | 9.996E-01 |  | x |
| ENSSSCG00000037015 | *SESN3* | -2.000 | -1.378 | -5.707 | 2.541E-07 | 2.477E-03 | x | x |
| ENSSSCG00000032452 | *WFS1* | -2.198 | -2.138 | -5.510 | 9.509E-03 | 9.996E-01 | x |  |
| ENSSSCG00000039548 | *PTGFR* | -1.634 | -1.590 | -4.915 | 8.804E-03 | 9.996E-01 |  | x |
| ENSSSCG00000013829 | *SYDE1* | -1.670 | -1.160 | -4.188 | 5.795E-04 | 6.000E-01 | x |  |
| ENSSSCG00000002265 | *FAM174B* | -1.244 | -1.726 | -4.179 | 5.385E-02 | 9.996E-01 |  | x |
| ENSSSCG00000016233 | *SERPINE2* | -1.735 | -2.060 | -3.603 | 5.684E-02 | 9.996E-01 | x | x |
| ENSSSCG00000006243 | *PENK* | -0.420 | -2.104 | -3.573 | 7.628E-01 | 9.996E-01 |  |  |
| ENSSSCG00000038879 | *RELB* | -1.272 | -1.056 | -3.512 | 3.659E-03 | 9.996E-01 | x |  |
| ENSSSCG00000023408 | *SAMD4A* | -1.328 | -1.156 | -3.509 | 4.486E-02 | 9.996E-01 |  | x |
| ENSSSCG00000008449 | *SLC3A1* | -1.014 | -1.154 | -3.491 | 5.859E-02 | 9.996E-01 |  |  |
| ENSSSCG00000014921 | *PRSS23* | -1.141 | -1.739 | -3.360 | 2.719E-01 | 9.996E-01 |  | x |
| ENSSSCG00000017186 | *RNF157* | -1.218 | -2.338 | -3.317 | 2.413E-01 | 9.996E-01 | x | x |
| ENSSSCG00000035403 | *RFX2* | -1.109 | -1.022 | -2.958 | 1.550E-01 | 9.996E-01 |  | x |
| ENSSSCG00000010893 |  | -0.655 | -1.352 | -2.931 | 4.068E-01 | 9.996E-01 |  | x |
| ENSSSCG00000031819 | *TP53I11* | -1.002 | -1.711 | -2.883 | 4.102E-01 | 9.996E-01 |  | x |
| ENSSSCG00000017137 | *METRNL* | -0.674 | -1.102 | -2.812 | 2.422E-01 | 9.996E-01 |  |  |
| ENSSSCG00000032562 | *TMC6* | -0.837 | -1.152 | -2.765 | 2.078E-01 | 9.996E-01 |  |  |
| ENSSSCG00000031261 | *RHOQ* | -0.903 | -1.046 | -2.750 | 1.839E-02 | 9.996E-01 |  |  |
| ENSSSCG00000001089 | *GPLD1* | -0.872 | -1.761 | -2.723 | 4.302E-01 | 9.996E-01 |  | x |
| ENSSSCG00000034259 | *PMEPA1* | -0.880 | -1.348 | -2.720 | 3.583E-01 | 9.996E-01 |  | x |
| ENSSSCG00000017014 | *PANK3* | -0.614 | -1.037 | -2.557 | 2.288E-01 | 9.996E-01 |  | x |
| ENSSSCG00000003377 | *ACOT7* | -0.790 | -2.688 | -2.544 | 3.439E-01 | 9.996E-01 | x |  |
| ENSSSCG00000010079 | *PPM1F* | -0.762 | -1.035 | -2.473 | 4.967E-02 | 9.996E-01 | x | x |
| ENSSSCG00000040464 | *LEP* | -0.747 | -2.186 | -2.463 | 1.880E-01 | 9.996E-01 | x | x |
| ENSSSCG00000022029 | *RAP1GAP* | -0.120 | -1.109 | -2.418 | 8.822E-01 | 9.996E-01 |  | x |
| ENSSSCG00000022099 | *TP53INP2* | -0.628 | -1.058 | -2.403 | 3.683E-01 | 9.996E-01 |  |  |
| ENSSSCG00000025652 | *CDH1* | -0.472 | -2.592 | -2.372 | 6.533E-01 | 9.996E-01 |  | x |
| ENSSSCG00000027266 | *PNPLA3* | -0.443 | -1.386 | -2.198 | 5.725E-01 | 9.996E-01 |  |  |
| ENSSSCG00000015401 | *PCLO* | -0.674 | -1.492 | -2.182 | 4.537E-01 | 9.996E-01 |  | x |
| ENSSSCG00000020872 |  | -1.029 | -1.128 | -2.090 | 1.340E-01 | 9.996E-01 |  |  |
| ENSSSCG00000032633 | *FAM53A* | -0.749 | -1.033 | -2.066 | 4.576E-02 | 9.996E-01 | x |  |
| ENSSSCG00000015559 | *NCF2* | -0.679 | -1.221 | -2.061 | 3.570E-01 | 9.996E-01 |  | x |
| ENSSSCG00000015766 | *WDR17* | -0.609 | -1.139 | -1.998 | 2.093E-01 | 9.996E-01 |  |  |
| ENSSSCG00000009761 | *NCOR2* | -0.681 | -1.421 | -1.913 | 2.643E-01 | 9.996E-01 |  |  |
| ENSSSCG00000016928 | *RAB3D* | -0.491 | -1.142 | -1.888 | 2.953E-01 | 9.996E-01 |  |  |
| ENSSSCG00000011230 | *OSBPL10* | -0.576 | -1.594 | -1.869 | 4.272E-01 | 9.996E-01 |  | x |
| ENSSSCG00000017298 | *TANC2* | -0.615 | -1.541 | -1.846 | 4.896E-01 | 9.996E-01 |  |  |
| ENSSSCG00000007899 |  | -0.524 | -1.036 | -1.814 | 4.500E-01 | 9.996E-01 |  | x |
| ENSSSCG00000026421 | *PKD2L2* | -0.463 | -1.230 | -1.800 | 5.098E-01 | 9.996E-01 |  |  |
| ENSSSCG00000015332 | *PON1* | -0.626 | -1.076 | -1.763 | 2.530E-01 | 9.996E-01 | x | x |
| ENSSSCG00000009215 | *ABCG2* | -0.455 | -1.446 | -1.749 | 5.684E-01 | 9.996E-01 |  |  |
| ENSSSCG00000017328 | *ARHGAP27* | -0.235 | -2.788 | -1.699 | 8.113E-01 | 9.996E-01 | x | x |
| ENSSSCG00000017199 | *TRIM47* | -0.362 | -1.057 | -1.645 | 6.717E-01 | 9.996E-01 |  | x |

ENSSSCG00000016928 did not have an annotated 3'-UTR and was therefore excluded from further analyses.
<sup>a</sup>Log<sub>2</sub>FC: estimated log<sub>2</sub> fold change for mean exonic fraction from adipocytes of *lean* and *obese* UNIK minipigs;
<sup>b</sup>ΔEx: exonic fraction increment (Ex<sub>2</sub> – Ex<sub>1</sub>) in log<sub>2</sub> scale when comparing exon abundances in obese (Ex<sub>1</sub>) vs lean
(Ex<sub>2</sub>) UNIK minipigs; <sup>c</sup>PTc: post-transcriptional signal (ΔEx – ΔInt) in z-score scale. The *q*-value has been calculated
with the false discovery rate (FDR) approach (Benjamini & Hochberg 1995); <sup>d</sup>DE: Significantly differentially
expressed (FC < -2; *q*-value < 0.05). The “x” symbols indicate significantly downregulated genes according to their
exonic counts, as well as those mRNA genes targeted by at least one of the significantly upregulated miRNAs
excluding redundant seeds (N = 4, **Table S10**).

**Figure 2:**
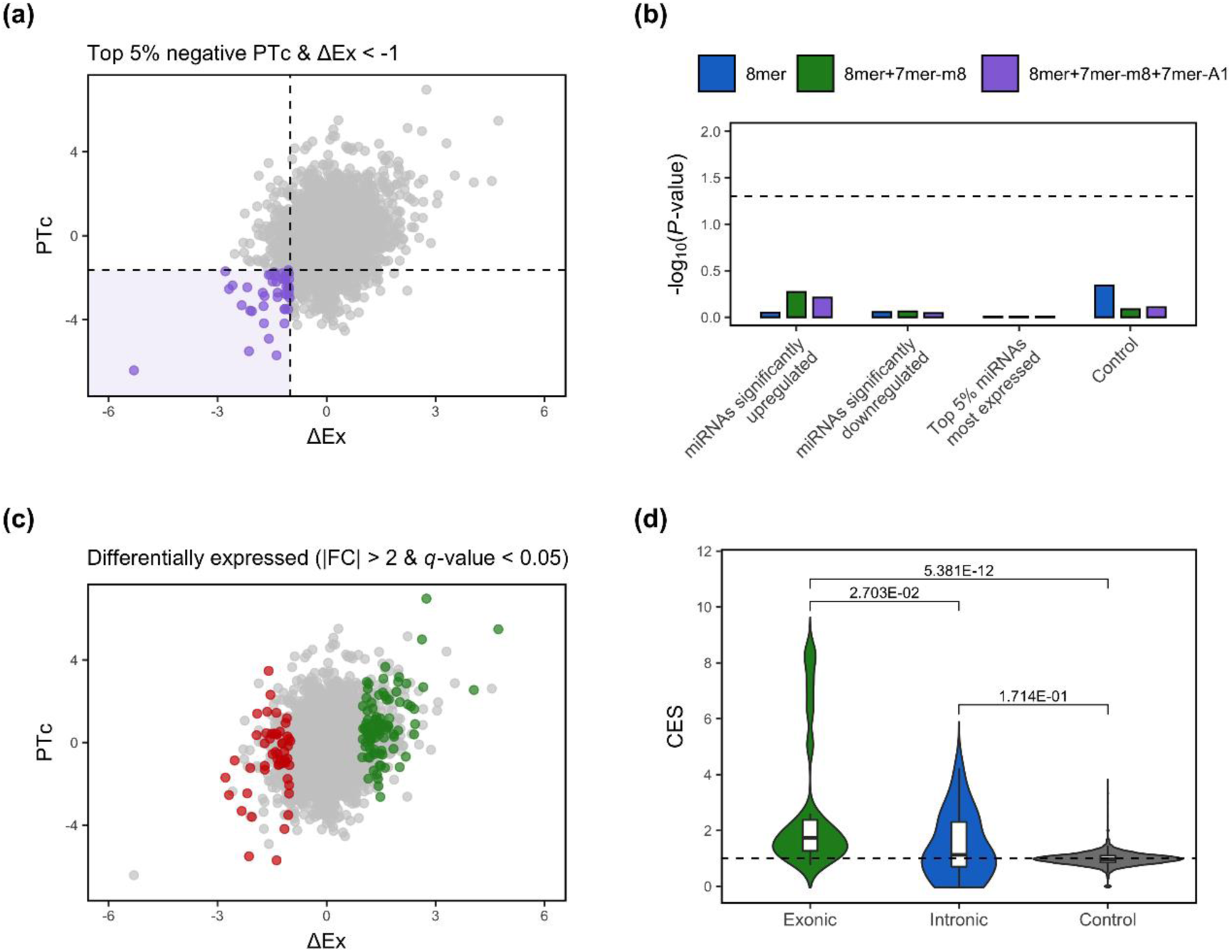
(**a**) Scatterplot depicting genes expressed in adipocytes obtained from UNIK minipigs with *lean* (N = 5) and *obese* (N = 5) phenotypes (using their body mass index as reference) according to their exonic fraction (ΔEx) and post-transcriptional (PTc) scores. Genes with the top 5% negative PTc scores and at least 2-fold ΔEx reduction (equivalent to -1 in the log_2_ scale) are highlighted in purple and delimited by dashed lines. (**b**) Enrichment analyses comparing all expressed mRNA genes and the set of mRNA genes with the top 5% negative PTc scores and at least 2-fold ΔEx reduction as being putatively targeted by either significantly upregulated miRNAs (FC > 1.5; *q*-value < 0.05), significantly downregulated miRNAs (FC < -1.5; *q*-value < 0.05) or the top 5% most highly expressed miRNAs, excluding significantly upregulated miRNAs. As indicated with the dashed line, a nominal *P*-value = 0.05 was set as a significance threshold. (**c**) Scatterplot depicting genes expressed in adipocytes obtained from UNIK minipigs with *lean* (N = 5) and *obese* (N = 5) phenotypes (using their body mass index as reference) according to their exonic fraction (ΔEx) and post-transcriptional (PTc) scores. Genes significantly upregulated are in green, while those being downregulated are in red (|FC| > 2; *q*-value < 0.05). (**d**) Covariation enrichment scores (CES) for the exonic and intronic fractions of the 25 mRNA genes with the top 5% negative PTc scores and at least 2-fold ΔEx reduction predicted to harbor binding sites for upregulated miRNAs (N = 4) in adipocytes obtained from UNIK minipigs with *lean* (N = 5) and *obese* (N = 5) phenotypes. The control set was established by generating 1,000 permuted lists of 25 genes chosen at random and using their exonic and intronic fractions for calculating their CES values. Statistical significance was assessed using a Mann-Whitney U non-parametric test (Mann & Whitney 1947). The dashed line represents a CES of 1, equivalent to an observed null fold change in covariation.

Twenty-five out of the 44 (58.14%, **Table S9b**) mRNAs downregulated in *lean* minipigs and displaying the top 5% negative PTc scores were classified as putative targets of the set of miRNAs upregulated in *lean* minipigs (N = 4, ssc-miR-92b-3p, ssc-miR-148a-3p, ssc-miR-204 and ssc-miR-214-3p; **Table S10** in bold). Target prediction and context-based pruning of miRNA-mRNA interactions for these 25 genes made possible to detect eight 8mer, 21 7mer-m8 and 24 7mer-A1 miRNA binding sites (**Table S12**) for upregulated miRNAs (N = 4) in *lean* UNIK minipigs (**Table S10**). Again, the *SESN3* gene showed the highest number of predicted putative miRNA target sites in its 3’-UTR (**Table S12**).

Enrichment analyses for these 25 mRNAs (**Table 2**) revealed no significant enrichment for 8mer, 7mer-m8 and 7mer-A1 miRNA binding sites (**Fig. 2b**). Besides, only seven of them (26.92%) appeared as significantly downregulated (FC < -2; *q*-value < 0.05) in the differential expression analyses with *edgeR* (**Table S9b** and **Table 2, Fig. 2c**). The exonic fractions of 18 out of these 25 mRNA genes showed significantly increased covariation (*P-*value = 2.703E-02) compared to the covariation observed for the intronic fractions (**Fig. 2d** and **Table S13**).

Regarding Tc scores, a total of 195 genes showed significant transcriptional signals (|FC| > 2; *q*-value < 0.05, **Table S11b**), and 48 of them were also significantly differentially expressed (24.61%, **Tables S9a** and **S11b**). Moreover, three of them (*ARHGAP27, CDH1* and *LEP*) were found among those with the top 5% post-transcriptional signals (**Table 2** and **Table S11b** in bold). The whole lists of expressed genes after EISA and their PTc and Tc scores are available at **Table S11c** and **S11d**, respectively.

Results obtained for qPCR verification analyses are described in **Fig. S5**.

## Discussion

### Contribution of the Tc and PTc components of gene regulation to energy homeostasis in porcine muscle and adipose tissues

After running EISA on both muscle and adipose tissue datasets, we observed that the number of genes with significant transcriptional signals (Tc) was much higher than that of loci with significant post-transcriptional signals (PTc). Such difference evidences that gene expression changes induced by feeding or obesity might be mostly driven by transcriptional rather than post-transcriptional modulators. It is worth noting, however, that few mRNAs showed post-transcriptional signals alone and were thus mixed with transcriptional signals either in concordant or in opposite directions.

For prioritizing putative post-transcriptionally repressed mRNA genes by miRNAs, we focused on those with the strongest observed downregulation based on their ΔEx values (at least 2-fold reduction) and PTc signal (top 5% negative scores). Hence, we did not consider the significance of PTc scores as a relevant criterion, as these will appear as significant only when the post-transcriptional response is strong and not confounded by a cooperative repressive transcriptional signal. Alternative thresholds other than the top 5% negative PTc scores or 2-fold for ΔEx fraction could be applied, depending on the strength of the post-transcriptional signal detected.

Besides, it is worth noting that our RNA-seq data was generated following manufacturer’s instructions for TruSeq stranded libraries. This commonly used protocol selects for poly(A) mRNAs, and the majority of non-poly(A) intronic lariats generated after splicing will be lost, thus producing an artificial decrease in the quantified intronic fraction (intronic reads). Although the intronic yield is decreased in poly(A) RNA compared to total RNA protocols (Ameur *et al*. 2011), the recoverable intronic fraction is still highly correlated with nascent transcription (Gaidatzis *et al*. 2015; La Manno *et al*. 2018), probably derived from the presence of poly(A) introns or unspliced mRNAs being sequenced, albeit at low abundance. Thus, the remaining intronic reads might conform a limited and indirect yet representative proxy of the transcriptional activity. Alternative methods for measuring such transcriptional activity by directly monitoring transcription across the genome have also been applied (Patel *et al*. 2020), and might be preferred over the use of intronic fractions. However, the implementation of such alternatives is limited so far, and the relatively simple and cost-effective usage of intronic reads present in already available RNA-seq data justifies the use of the EISA approach.

### Differential expression analyses and EISA highlight different sets of genes

Few genes with significant Tc and PTc components were also classified by *edgeR* as significantly differentially expressed. Such a discrepancy between EISA and differential expression analyses is in agreement with the subtle regulation elicited by miRNAs, which is dependent on the expression level of miRNAs and the number of target sites within a given mRNA 3’-UTR (Bartel 2018). In this way, EISA might serve as a good approach to identify both strong and subtle post-transcriptional effects mediated by miRNAs that canonical differential expression approaches might not be able to capture. Importantly, these discrepancies were reduced when we focused on genes with top 5% negative PTc scores and at least 2-fold reduction in their ΔEx values: as much as 69.23% (skeletal muscle) and 27.27% (adipose tissue) of such genes were also detected as differentially expressed. This increase in concordance was more pronounced in the skeletal muscle (fasted vs fed gilts) experimental system. This might be due to the overall stronger upregulation of miRNAs observed for this dataset when compared with that generated in the adipose tissue experiment, which can be explained by intrinsic genomic differences among pig breeds, the tissues analyzed and/or the metabolic challenge undertaken.

### Predicting the contribution of miRNAs to the post-transcriptional regulatory response in porcine muscle and adipose tissues

Since the efficacy of miRNA-based repression of mRNA expression depends on the context of the miRNA binding site within the 3’-UTR (Grimson *et al*. 2007), we have assessed the usefulness of introducing context-based filtering criteria for removing spurious *in silico*-predicted binding sites for miRNAs. Using enrichment analyses, we were able to link the expression of the set of mRNAs with downregulated exonic fraction to the expression of upregulated miRNAs predicted to target them.

In the skeletal muscle system, prediction of miRNA binding sites in mRNA genes displaying the top 5% negative post-transcriptional signals and at least 2-fold reduction in their exonic fraction revealed that the majority of them (80.77%) harbored at least one binding site for the corresponding set of significantly upregulated miRNAs (N = 6). In contrast, such pattern was much less evident in the adipose tissue, which could be explained by the fact that the majority of upregulated miRNAs in *lean* minipigs did not reach significance.

Although we verified by qPCR the RNA-seq expression levels of selected downregulated mRNAs and upregulated miRNAs in the UNIK adipose tissue experiment, further experimental validation of the reported mRNA-miRNA interactions is needed. *In silico* predictions of miRNA binding sites, EISA and covariation analyses might be helpful to identify miRNA-mRNA pairs to be experimentally validated with co-transfection gene reporter assays, in order to increase the yet scarce collection of validated mRNA-miRNA interactions in domestic species.

### Covariation patterns in the expression of downregulated mRNAs predicted to be targeted by upregulated microRNAs

We further hypothesized that mRNA genes showing relevant post-transcriptional downregulation might be repressed by the same set of significantly upregulated miRNAs, which could induce shared covariation in the expression profiles of such mRNAs at the exonic level. In contrast, their intronic fraction would be mainly unaffected as introns would have been excised prior to any given miRNA-driven downregulation. In this way, an increased gene covariation in downregulated mRNAs with high post-transcriptional signals might be detectable at the exon but not at the intron level. Indeed, our results revealed an increased covariation in downregulated mRNAs with high post-transcriptional signals at their exonic fraction compared with covariation patterns of their intronic fraction, suggesting that their expression might be repressed by a common set of upregulated miRNAs.

### Genes displaying the strongest post-transcriptional signals in porcine skeletal muscle and adipose tissue are involved in glucose and lipid metabolism

The mRNA genes showing the strongest post-transcriptional downregulation in fasted vs fed gilts displayed a variety of relevant biological functions. The *DKK2* gene showed the most negative significant PTc score. This gene also displayed the strongest covariation difference in its exonic fraction compared with the intronic one. This consistent post-transcriptional regulatory effect might be mediated by ssc-miR-421-5p and ssc-miR-30a-3p, two highly significantly upregulated miRNAs. The DKK2 protein is a member of the dickkopf family, which inhibits the Wnt signaling pathway through its interaction with the LDL-receptor related protein 6 (LRP6). Its repression has been associated with reduced blood-glucose levels and improved glucose uptake (Li *et al*. 2012), as well as with improved adipogenesis (Yang & Shi 2021) and the inhibition of aerobic glycolysis (Deng *et al*. 2019). These results are consistent with the increased glucose usage and triggered adipogenesis in muscle tissue after nutrient supply. Other relevant post-transcriptionally downregulated mRNAs detected with EISA were: pyruvate dehydrogenase kinase 4 (*PDK4*), interleukin 18 (*IL18*), nuclear receptor subfamily 4 group A member 3 (*NR4A3*), acetylcholine receptor subunit α (*CHRNA1*), PBX homeobox 1 (*PBX1*), Tet methylcytosine dioxygenase 2 (*TET2*), BTB domain and CNC homolog (*BACH2*), all of which are involved in the regulation of energy homeostasis (Lindegaard *et al*. 2013; Zhang *et al*. 2014; Tamahara *et al*. 2017; Wu *et al*. 2018; Xu *et al*. 2019) and lipid metabolism in muscle cells (Monteiro *et al*. 2011; Pearen *et al*. 2013) in response to nutrient uptake.

On the other hand, several genes with high post-transcriptional signals were not predicted to be targets of upregulated miRNAs: the circadian associated repressor of transcription (*CIART*) and period 1 (*PER1*), oxysterol binding protein like 6 (*OSBPL6*) and nuclear receptor subfamily 4 group A member 1 (*NR4A1*). Interestingly, all of them were significantly downregulated (Cardoso *et al*. 2017; Mármol-Sánchez *et al*. 2020). An explanation to this might be that although miRNAs are key post-transcriptional regulators, other alternative post-transcriptional effectors, such as long non-coding RNAs (lncRNAs), circular RNAs (circRNAs) or RNA binding proteins (RBPs) might be at play. Besides, indirect repression via upregulated miRNAs acting over regulators of these genes, such as transcription factors, could be also a major influence on their observed repression (Patel *et al*. 2020).

The use of EISA on expression data from adipocytes isolated from *obese* vs *lean* UNIK minipigs revealed several mRNA genes with high post-transcriptional repression, which are also involved in the regulation of lipid metabolism and energy homeostasis. The gene showing the highest post-transcriptional signal was the estrogen related receptor γ (*ESRRG*), which modulates oxidative metabolism and mitochondrial function in adipose tissue and inhibits adipocyte differentiation when repressed (Kubo *et al*. 2009). Another relevant locus identified with EISA was *SESN3*, an activator of the mTORC2 and PI3K/AKT signaling pathway that promotes lipolysis when inhibited (Tao *et al*. 2015). This gene showed the most significant downregulation in *lean* minipigs, and gathered multiple putative binding sites for all the four upregulated miRNAs under study.

Other genes showing significant post-transcriptional downregulation were: sterile α motif domain containing 4A (*SAMD4A*), prostaglandin F2-receptor protein (PTGFR), serine protease 23 (PRSS23), ring finger protein 157 (*RNF157*), oxysterol binding protein like 10 (*OSBPL10*), glycosylphosphatidylinositol phospholipase 1 (GPLD1), RAP1 GTPase activating protein (RAP1GAP) and leptin (*LEP*), and all of which are tightly linked to adipocyte differentiation (Perttilä *et al*. 2009; Martínez *et al*. 2013; Chen *et al*. 2014; Kosacka *et al*. 2018) or energy homeostasis (Ussar *et al*. 2012; Wang *et al*. 2018; Izquierdo *et al*. 2019; Kuo *et al*. 2020). Despite the overall weak influence of putative miRNA-driven downregulation on mRNAs expressed in adipocytes, we were able to identify a set of genes with high post-transcriptional signals indicative of putative miRNA-derived repression and tightly related to adipose tissue metabolism regulation. However, non-miRNA transcriptional and post-transcriptional modulators, might also contribute to such repression.

## Conclusions

EISA applied to study gene regulation in porcine skeletal muscle and adipose tissues showed that more genes were subjected to transcriptional rather than post-transcriptional regulation, suggesting that changes in mRNA levels are mostly driven by factors acting at the transcriptional level. More importantly, the concordance between the sets of significantly differentially expressed genes and those with significant Tc or PTc scores was quite limited, but improved (mostly in the skeletal muscle experiment) when we prioritized the downregulated genes with the top 5% negative post-transcriptional signals. Nevertheless, many of the genes with relevant PTc signals were not among the top significantly downregulated loci, thus demonstrating the usefulness of complementing differential expression analyses with the EISA approach. In the skeletal muscle, we detected several mRNAs predicted to be co-regulated by a common set of miRNAs. In contrast, in the adipose tissue such relationship was more subtle, suggesting that the contribution of miRNAs to mRNA repression might be affected by tissue type, breed and/or intrinsic experimental factors. Finally, EISA made possible to identify several genes related with carbohydrate and lipid metabolism, which may play relevant roles in the energy homeostasis of the skeletal muscle and adipose tissues. By differentiating the transcriptional from the post-transcriptional changes in mRNA expression, EISA provides a valuable view, complementary to differential expression analysis, about the miRNA-driven regulation of gene expression.

## Ethics approval

Animal care and management procedures for Duroc gilts followed national guidelines for the Good Experimental Practices and were approved by the Ethical Committee of the Institut de Recerca i Tecnologia Agroalimentàries (IRTA). Animal care and management procedures for UNIK minipigs were carried out according to the Danish “Animal Maintenance Act” (Act 432 dated 9 June 2004).

## Availability of data

The RNA-seq and small RNA-seq datasets from skeletal muscle tissue used in the current study are available at the Sequence Read Archive (SRA) database with BioProject codes PRJNA386796 and PRJNA595998, respectively. For the adipose tissue samples, RNA-seq and small RNA-seq datasets are available at PRJNA563583 and PRJNA759240.

## Competing interests

The authors declare that they have no competing interests

## Funding

The present research work was funded by grants AGL2013-48742-C2-1-R and AGL2013-48742-C2-2-R awarded by the Spanish Ministry of Economy and Competitivity. E. Mármol-Sánchez was funded with a PhD fellowship FPU15/01733 awarded by the Spanish Ministry of Education and Culture (MECD). YRC is recipient of a Ramon y Cajal fellowship (RYC2019-027244-I) from the Spanish Ministry of Science and Innovation.

## Authors’ contributions

The authors’ responsibilities were as follows: M.A. and R.Q. generated the total RNA and small RNA sequencing data corresponding to the GM muscle, respectively. S.C., M.J.J., C.B.J. and M.F produced the total RNA sequencing corresponding to the adipose tissue. M.A., R.Q., S.C., M.J.J., C.B.J., M.F., T.F.C. and E.M.-S. conducted the research. S.C. performed qPCR analyses. E.M.-S. analyzed the data. L.M.Z. contributed to bioinformatic analyses. Y.R.-C. contributed to critical assessment. M.A. and R.Q. secured funding for the study. E.M.-S., S.C. and M.A. drafted the manuscript. All authors contributed to manuscript corrections, read and approved the final manuscript.

## Acknowledgements

The authors would like to thank the Department of Veterinary Animal Sciences in the Faculty of Health and Medical Sciences of the University of Copenhagen for providing sequencing data and their facilities and resources for qPCR experiments. We also acknowledge Selección Batallé S.A. for providing animal material and the support of the Spanish Ministry of Economy and Competitivity for the Center of Excellence Severo Ochoa 2020–2023 (CEX2019-000902-S) grant awarded to the Centre for Research in Agricultural Genomics (CRAG, Bellaterra, Spain). Thanks also to the CERCA Programme of the Generalitat de Catalunya for their support.

## Supplementary Materials

**Figure S1:** Diagram depicting the routine/pipeline implemented for studying miRNA-driven post-transcriptional regulatory signals applying the EISA approach and additional enrichment and covariation analyses.

**Figure S2:** Diagram representing each one of the context-based filtering criteria used for excluding *in silico*-predicted miRNA-mRNA interactions. AU: miRNA binding sites with AU-rich flanking sequences (30 nts upstream and downstream). M: miRNA binding sites located in the middle of the 3’-UTR sequence (45-55%). E: miRNA binding sites located too close (< 15 nts) to the beginning or the end of the 3’-UTR sequences.

**Figure S3:** Scatterplots depicting the exonic (ΔEx) and intronic (ΔInt) fractions of expressed genes from *gluteus medius* skeletal muscle samples of fasting (*AL-T0*, N = 11) and fed (*AL-T2*, N = 12) Duroc gilts. (**a**) Genes differentially expressed and showing either significant upregulation (FC > 2; *q*-value < 0.05, in green) or downregulation (FC < -2, *q*-value < 0.05, in red) in fed (*AL-T2*, N = 12) Duroc gilts with respect to their fasted (*AL-T0*, N = 11) counterparts. (**b**) Genes with the top 5% post-transcriptional (PTc) negative scores and at least 2-fold reduced exonic (ΔEx) fraction (equivalent to -1 in the log_2_ scale) are highlighted in purple.

**Figure S4:** Enrichment analyses comparing all expressed mRNA genes and the set of mRNA genes with the (**a**) top 1% and (**b**) top 5% negative PTc scores and at least 2-fold ΔEx reduction as being putatively targeted by significantly upregulated miRNAs (FC > 1.5; *q*-value < 0.05) from *gluteus medius* skeletal muscle samples of fasting (*AL-T0*, N = 11) and fed (*AL-T2*, N = 12) Duroc gilts. Results show the change in enrichment significance (expressed as -log_10_ of the estimated *P*-value) when incorporating context-based pruning of 8mer, 7mer-m8 and 7mer-A1 miRNA binding sites. R: Raw enrichment analyses without any additional context-based pruning. AU: Enrichment analyses removing miRNA binding sites without AU-rich flanking sequences (30 nts upstream and downstream). M: Enrichment analyses removing miRNA binding sites located in the middle of the 3’-UTR sequence (45-55%). E: Enrichment analyses removing miRNA binding sites located too close (< 15 nts) to the beginning or the end of the 3’-UTR sequences. The dashed line represents a nominal *P*-value of 0.05 set as significance threshold.

**Figure S5:** Quantification of selected genes and miRNAs expressed in the pig adipose tissue by qPCR. (**a**) Barplots depicting qPCR log_2_ transformed relative quantities (Rq) for *LEP, OSBPL10, PRSS23, RNF157* and *SERPINE2* mRNA transcripts measured in adipocytes from the retroperitoneal fat of *lean* (N = 5) and *obese* (N = 5) UNIK minipigs. (**b**) Barplots depicting qPCR log_2_ transformed relative quantities (Rq) for ssc-miR-148a-3p, ssc-miR-214-3p and ssc-miR-92b-3p miRNA transcripts measured in isolated adipocytes from the retroperitoneal fat of *lean* (N = 5) and *obese* (N = 5) UNIK minipigs. All the analyzed mRNA genes showed a reduced expression in *lean* minipigs compared with their *obese* counterparts, and the *LEP* gene was the most significantly downregulated gene. For miRNAs, the opposite pattern of expression was observed, being all of them upregulated in *lean* minipigs. Moreover, ssc-miR-92b-3p showed the most significant increased expression in *lean* minipigs, in agreement with results obtained in differential expression analyses (**Table S10**).

**Table S1:** Phenotypic values of body mass index (BMI) trait and sex classification recorded in 11 Duroc-Göttingen minipigs from the F2-UNIK resource population and grouped as *obese* or *lean* in accordance with their BMI.

**Table S2:** Primers for qPCR verification of selected mRNAs and miRNAs in the F2-UNIK Duroc-Göttingen minipig population after comparing *lean* (N = 5) and *obese* (N = 5) individuals.

**Table S3:** Raw Cq values obtained in qPCR analyses measuring adipocyte expression levels of selected mRNAs and miRNAs from *lean* (N = 5) and *obese* (N = 5) UNIK minipigs.

**Table S4:** Differential expression analyses of RNA-seq data using the *edgeR* tool and comparing *gluteus medius* expression profiles of fasted *AL-T0* (N = 11) and fed *AL-T2* (N = 12) Duroc gilts. (**a**) Differentially expressed genes (q-value < 0.05, N = 454). In bold are genes either upregulated (N = 52) or downregulated (N = 80) with |FC| > 2 and q-value < 0.05. (**b**) Differential expression results for genes with top 5% post-transcriptional (PTc) scores and at least 2-fold reduced exonic fraction (ΔEx) (equivalent to -1 in the log_2_ scale). The 18 genes with FC < -2 and *q*-value < 0.05 are shown in bold. (**c**) Differential expression results showing the whole list of expressed genes with an average expression above 1 CPM in at least 50% of samples within each group (N = 9,492).

**Table S5:** Differential expression analyses of microRNAs using the *edgeR* tool and comparing *gluteus medius* expression levels of fasted *AL-T0* (N = 11) and fed *AL-T2* (N = 12) Duroc gilts. In bold are microRNAs detected as significantly differentially expressed (|FC| > 1.5; *q*-value < 0.05, N = 16).

**Table S6:** Post-transcriptional (PTc) and transcriptional (Tc) signals detected with EISA in genes expressed in *gluteus medius* skeletal muscle samples from fasted (*AL-T0*, N = 11) and fed (*AL-T2*, N = 12) Duroc gilts. (**a**) Genes with significant PTc scores (|FC| > 2; *q*-value < 0.05, N = 133). In bold are genes with at least 2-fold reduced ΔEx fraction (N = 3). (**b**) Genes with significant Tc scores (|FC| > 2; *q*-value < 0.05, N = 344). In bold are genes also significant in their PTc scores (|FC| > 2; *q*-value < 0.05, N = 91). EISA results showing the whole list of genes with an average expression above 1 CPM in at least 50% of samples within each group (N = 9,492) and their (**c**) PTc and (**d**) Tc scores.

**Table S7:** Binding sites in the 3’-UTRs of mRNA genes (with the top 5% negative PTc scores and at least 2-fold reduction in the exonic fraction) predicted as targets (N = 21) of non-redundant significantly upregulated miRNAs (N = 6) expressed in the *gluteus medius* skeletal muscle samples from fasting (*AL-T0*, N = 11) and fed (*AL-T2*, N = 12) Duroc gilts.

**Table S8:** Covariation enrichment scores (CES) for the exonic and intronic fractions of mRNA genes (with the top 5% negative post-transcriptional signals PTc and at least 2-fold reduction in their exonic ΔEx fraction) predicted as targets of non-redundant significantly upregulated miRNAs (N = 6) expressed in *gluteus medius* skeletal muscle samples from fasting (*AL-T0*, N = 11) and fed (*AL-T2*, N = 12) Duroc gilts.

**Table S9:** Differential expression analyses of RNA-seq data using the *edgeR* tool and comparing adipocyte expression profiles of *lean* (N = 5) and *obese* (N = 5) UNIK minipigs classified in accordance with their body mass index. (**a**) Differentially expressed genes (*q*-value < 0.05, N = 299). In bold are genes either upregulated (N = 52) or downregulated (N = 95) with |FC| > 2 and *q*-value < 0.05. (**b**) Differential expression results for genes with top 5% post-transcriptional (PTc) scores and at least 2-fold reduced exonic fraction (ΔEx) (equivalent to -1 in the log_2_ scale). The 12 genes with FC < -2 and *q*-value < 0.05 are shown in bold. (**c**) Differential expression results showing the whole list of expressed genes with an average expression above 1 CPM in at least 50% of samples within each group (N = 9,746).

**Table S10:** Differential expression analyses of microRNAs using the *edgeR* tool and comparing adipocyte expression profiles from *lean* (N = 5) and *obese* (N = 5) UNIK minipigs classified in accordance with their body mass index. In bold are microRNAs detected as significantly differentially expressed (|FC| > 1.5; *P*-value < 0.01, N = 7).

**Table S11:** Post-transcriptional (PTc) and transcriptional (Tc) signals detected with EISA in genes expressed in adipocytes from *lean* (N = 5) and *obese* (N = 5) UNIK minipigs classified in accordance with their body mass index. (**a**) Genes with significant PTc scores (|FC| > 2; *q*-value < 0.05, N = 1). In bold are genes with at least 2-fold reduced ΔEx fractions (N = 1). (**b**) Genes with significant Tc scores (|FC| > 2; *q*-value < 0.05, N = 195). In bold are genes also among the top 5% negative PTc scores and at least 2-fold ΔEx reduction (N = 3). EISA results showing the whole list of genes with an average expression above 1 CPM in at least 50% of samples within each group (N = 9,746) and their (**c**) PTc and (**d**) Tc scores.

**Table S12:** Binding sites in the 3’-UTRs of mRNA genes (with the top 5% negative PTc scores and at least 2-fold reduction in the exonic fraction) predicted as targets (N = 25) of non-redundant significantly upregulated miRNAs (N = 4) expressed in adipocytes from *lean* (N = 5) and *obese* (N = 5) UNIK minipigs.

**Table S13:** Covariation enrichment scores (CES) for the exonic and intronic fractions of mRNA genes (with the top 5% negative post-transcriptional signals PTc and at least 2-fold reduction in their exonic ΔEx fraction) predicted as targets of non-redundant significantly upregulated miRNAs (N = 4) expressed in adipocytes from *lean* (N = 5) and *obese* (N = 5) UNIK minipigs.

## Notes

### Competing Interest Statement

The authors have declared no competing interest.

### Summary of Updates

A corrected and reduced version after co-author revision.

