## Supplementary material for "Modeling microRNA-driven post-transcriptional regulation by using exon-intron split analysis (EISA) in pigs": Suppl Figures

[1]Centre for Research in Agricultural Genomics (CRAG), CSIC-IRTA-UAB-UB, Universitat Autònoma de Barcelona, 08193 Bellaterra, Spain. [2]Department of Veterinary and Animal Sciences, Faculty of Health and Medical Sciences, University of Copenhagen, 1871 Frederiksberg C, Denmark. [3]Universidad Nacional de Villa María, Villa María, Córdoba, Argentina. [4]Animal Breeding and Genetics Program, Institute for Research and Technology in Food and Agriculture (IRTA), Torre Marimon, 08140 Caldes de Montbui, Barcelona, Spain. [5]Departament de Ciència Animal i dels Aliments, Universitat Autònoma de Barcelona, 08193 Bellaterra, Barcelona, Spain.

*Emilio Mármol-Sánchez current affiliation: [1]Department of Molecular Biosciences, The Wenner-Gren Institute, Stockholm University, Stockholm, Sweden. [2]Centre for Paleogenetics, Stockholm University, Stockholm, Sweden.

[†]Tainã Figueiredo Cardoso current address: Embrapa Pecuária Sudeste, Empresa Brasileira de Pesquisa Agropecuária (EMBRAPA), 13560-970, São Carlos, SP, Brazil.

**Corresponding author:** Emilio Mármol-Sánchez. Science for Life Laboratory, Department of Molecular Biosciences, The Wenner-Gren Institute. Stockholm University, Stockholm, Sweden.

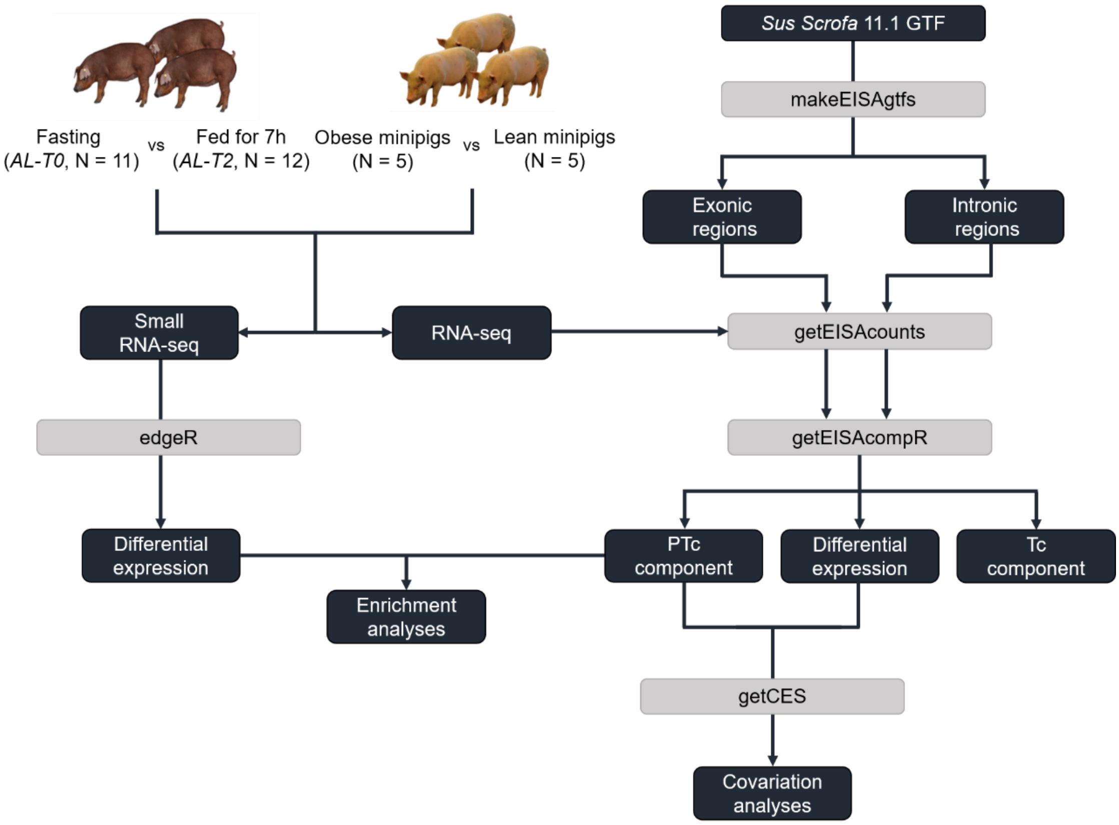


**Figure S1:** Diagram depicting the routine/pipeline implemented for studying miRNA-driven post-transcriptional regulatory signals applying the EISA approach and additional enrichment and covariation analyses.

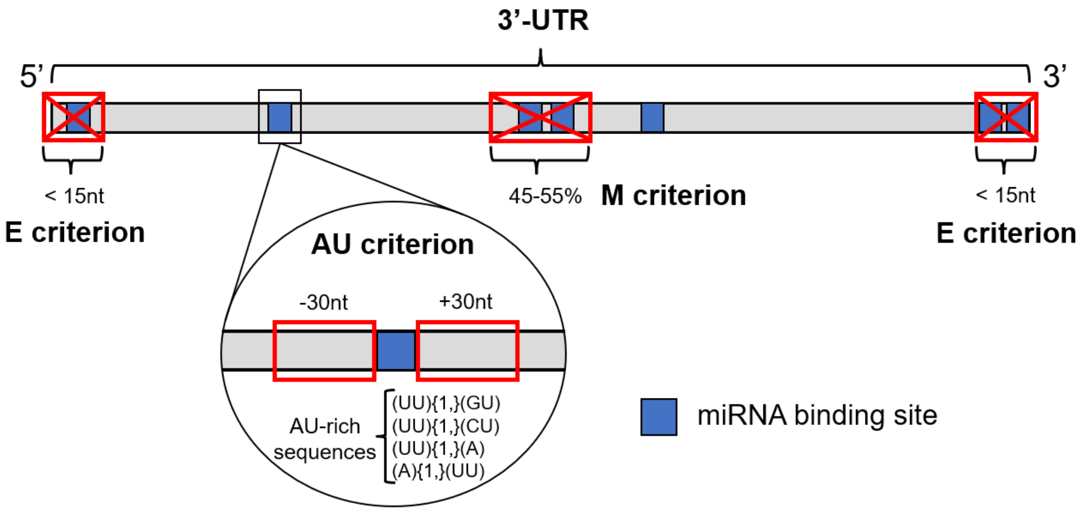


**Figure S2:** Diagram representing each one of the context-based filtering criteria used for excluding *in silico*-predicted miRNA-mRNA interactions. AU: miRNA binding sites with AU-rich flanking sequences (30 nts upstream and downstream). M: miRNA binding sites located in the middle of the 3'-UTR sequence (45-55%). E: miRNA binding sites located too close (< 15 nts) to the beginning or the end of the 3'-UTR sequences.

**(a)**

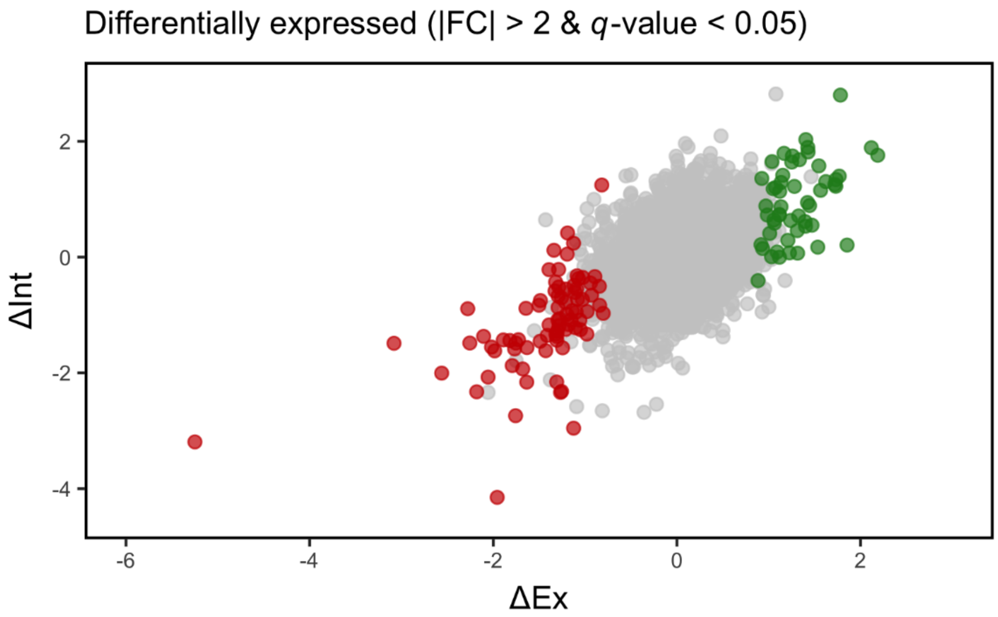
Differentially expressed (|FC| > 2 & *q*-value < 0.05)
ΔInt
2
0
-2
-4
-6
-4
-2
0
2
ΔEx

**(b)**

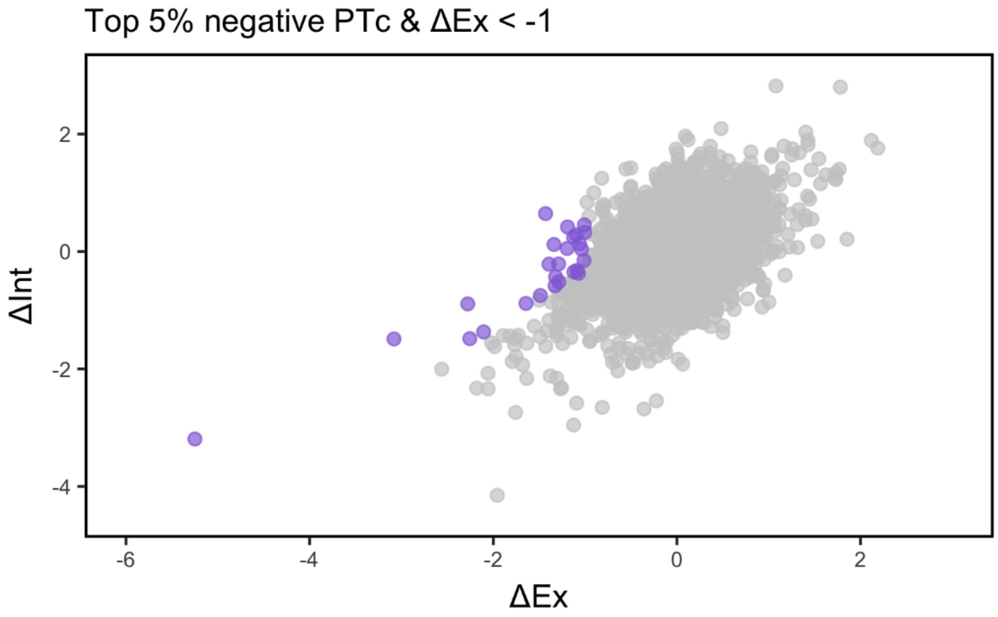
Top 5% negative PTc & ΔEx < -1
ΔInt
2
0
-2
-4
-6
-4
-2
0
2
ΔEx

**Figure S3:** Scatterplots depicting the exonic (ΔEx) and intronic (ΔInt) fractions of expressed genes from *gluteus medius* skeletal muscle samples of fasting (*AL-T0*, N = 11) and fed (*AL-T2*, N = 12) Duroc gilts. (**a**) Genes differentially expressed and showing either significant upregulation (FC > 2; *q*-value < 0.05, in green) or downregulation (FC < -2, *q*-value < 0.05, in red) in fed (*AL-T2*, N = 12) Duroc gilts with respect to their fasted (*AL-T0*, N = 11) counterparts. (**b**) Genes with the top 5% post-transcriptional (PTc) negative scores and at least 2-fold reduced exonic (ΔEx) fraction (equivalent to -1 in the $\log_2$ scale) are highlighted in purple.

**(a)**

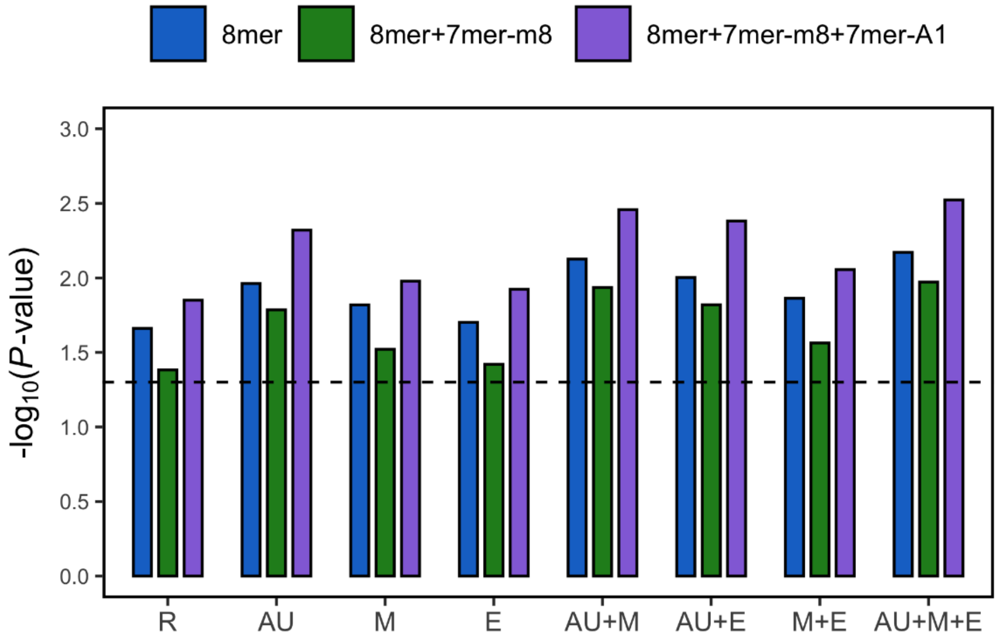

8mer
8mer+7mer-m8
8mer+7mer-m8+7mer-A1
-log10(P-value)
3.0
2.5
2.0
1.5
1.0
0.5
0.0
R
AU
M
E
AU+M
AU+E
M+E
AU+M+E


**(b)**

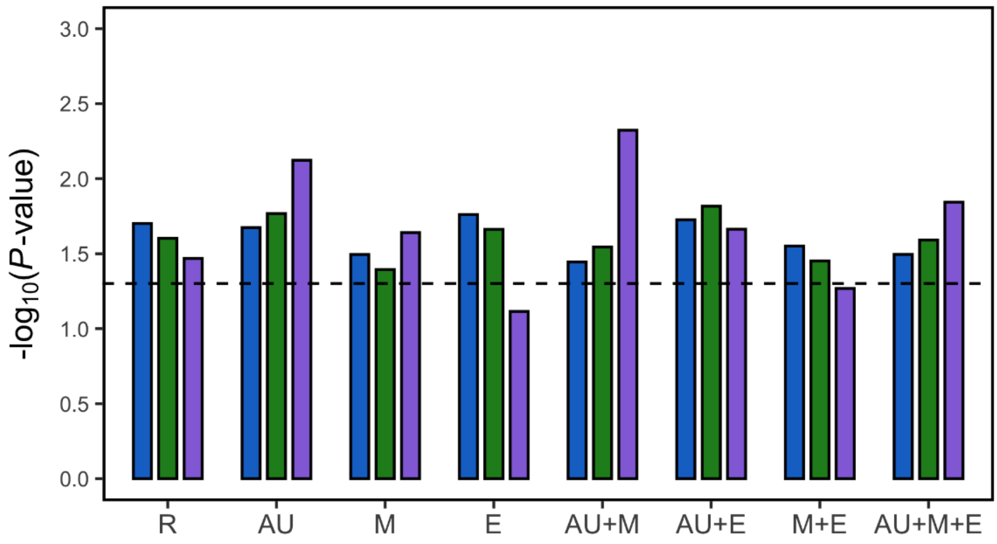

-log10(P-value)
3.0
2.5
2.0
1.5
1.0
0.5
0.0
R
AU
M
E
AU+M
AU+E
M+E
AU+M+E

**Figure S4:** Enrichment analyses comparing all expressed mRNA genes and the set of mRNA genes with the (**a**) top 1% and (**b**) top 5% negative PTc scores and at least 2-fold $\Delta$Ex reduction as being putatively targeted by significantly upregulated miRNAs (FC > 1.5; *q*-value < 0.05) from *gluteus medius* skeletal muscle samples of fasting (*AL-T0*, N = 11) and fed (*AL-T2*, N = 12) Duroc gilts. Results show the change in enrichment significance (expressed as $-\log_{10}$ of the estimated *P*-value) when incorporating context-based pruning of 8mer, 7mer-m8 and 7mer-A1 miRNA binding sites. R: Raw enrichment analyses without any additional context-based pruning. AU: Enrichment analyses removing miRNA binding sites without AU-rich flanking sequences (30 nts upstream and downstream). M: Enrichment analyses removing miRNA binding sites located in the middle of the 3'-UTR sequence (45-55%). E: Enrichment analyses removing miRNA binding sites located too close (< 15 nts) to the beginning or the end of the 3'-UTR sequences. The dashed line represents a nominal *P*-value of 0.05 set as significance threshold.

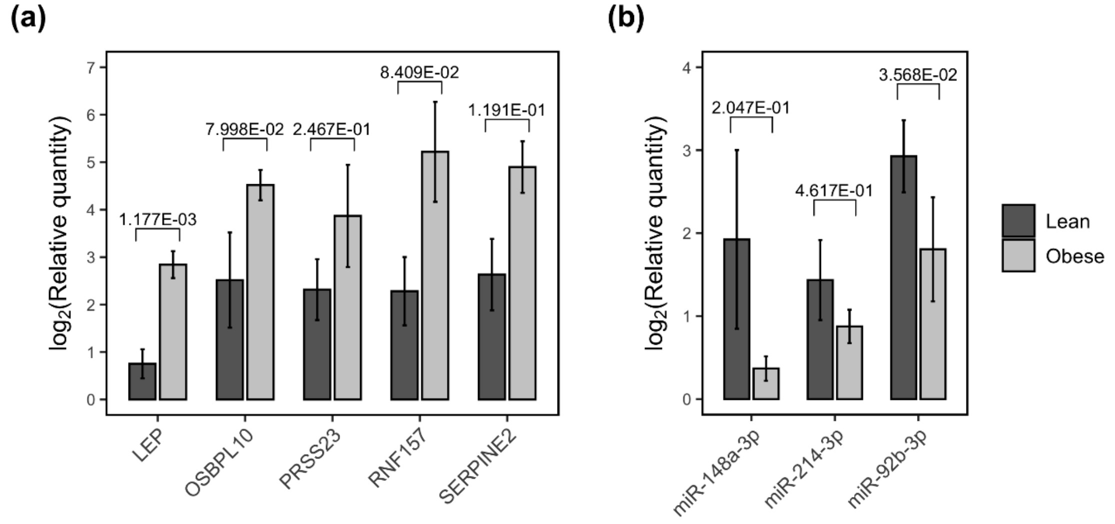


**Figure S5:** Quantification of selected genes and miRNAs expressed in the pig adipose tissue by qPCR. (**a**) Barplots depicting qPCR $log_2$ transformed relative quantities (Rq) for *LEP*, *OSBPL10*, *PRSS23*, *RNF157* and *SERPINE2* mRNA transcripts measured in adipocytes from the retroperitoneal fat of *lean* (N = 5) and *obese* (N = 5) UNIK minipigs. (**b**) Barplots depicting qPCR $log_2$ transformed relative quantities (Rq) for ssc-miR-148a-3p, ssc-miR-214-3p and ssc-miR-92b-3p miRNA transcripts measured in isolated adipocytes from the retroperitoneal fat of *lean* (N = 5) and *obese* (N = 5) UNIK minipigs. All the analyzed mRNA genes showed a reduced expression in *lean* pigs compared with their *obese* counterparts, and the *LEP* gene was the most significantly downregulated gene. For miRNAs, the opposite pattern of expression was observed, being all of them upregulated in *lean* minipigs. Moreover, ssc-miR-92b-3p showed the most significant increased expression in *lean* minipigs, in agreement with results obtained in differential expression analyses (**Table S10**).
