## Supplementary material for "Modeling microRNA-driven post-transcriptional regulation by using exon-intron split analysis (EISA) in pigs": Suppl Methods

[1]Centre for Research in Agricultural Genomics (CRAG), CSIC-IRTA-UAB-UB, Universitat Autònoma de Barcelona, 08193 Bellaterra, Spain. [2]Department of Veterinary and Animal Sciences, Faculty of Health and Medical Sciences, University of Copenhagen, 1871 Frederiksberg C, Denmark. [3]Universidad Nacional de Villa María, Villa María, Córdoba, Argentina. [4]Animal Breeding and Genetics Program, Institute for Research and Technology in Food and Agriculture (IRTA), Torre Marimon, 08140 Caldes de Montbui, Barcelona, Spain. [5]Departament de Ciència Animal i dels Aliments, Universitat Autònoma de Barcelona, 08193 Bellaterra, Barcelona, Spain.

*Emilio Mármol-Sánchez current affiliation: [1]Department of Molecular Biosciences, The Wenner-Gren Institute, Stockholm University, Stockholm, Sweden. [2]Centre for Paleogenetics, Stockholm University, Stockholm, Sweden.

[†]Tainã Figueiredo Cardoso current affiliation: Embrapa Pecuária Sudeste, Empresa Brasileira de Pesquisa Agropecuária (EMBRAPA), 13560-970, São Carlos, SP, Brazil.

**Corresponding author:** Emilio Mármol-Sánchez. Science for Life Laboratory, Department of Molecular Biosciences, The Wenner-Gren Institute. Stockholm University, Stockholm, Sweden.

**Covariation enrichment score (CES)**

The CES value for a given gene set reflects the frequency with which the mRNA expression correlation between members of that set is significant relative to the correlation between members of the whole expression data set. If multiple genes are downregulated by any upregulated miRNAs in a coordinated manner, we would expect to observe a reduced abundance in their mature spliced mRNA forms but not in the corresponding primary transcripts, i.e. we would detect a covariation only for their exonic fractions (but not for the intronic ones). In other words, the intronic fraction, eventually spliced and degraded in the nucleus, should not reflect any posterior post-transcriptional regulatory effects in the cytoplasm, so little or null covariation might be expected.

In this way, we can represent the variability in covariation within a set of genes as a fold change comparing the increase or reduction of observed significant covariation events relative to the remaining genes in the correlation matrix, and computed as follows:

Being $M$ the pairwise correlation square matrix obtained when computing Spearman´s correlation coefficients ($\rho$) from exonic and intronic fractions from the whole set of expressed mRNA genes with $q$-value < 0.05 after differential expression analyses and including any gene of interest (i.e. those with $q$-value > 0.05 and targeted by upregulated miRNAs and with the top 5% negative post-transcriptional signals and a reduction of at least 2-fold in their exonic fractions not previously included, in our case, **Table S4b** and **S9b**), i.e. $M = \{m_{ij}\}$, each pairwise correlation values are set to 1 if the correlation between a given pair of genes {i, j} is > |0.6| and significant according to the PCIT algorithm (Reverter & Chan 2008; Watson-Haigh *et al*. 2010), or set to zero otherwise. Then, we define $D = M[i_B, j_B]$ as the submatrix corresponding to the set genes of interest, with $i, j_B = \{\cup_{index_S}\}$, i.e. $D$ is a square matrix of size $d$, being $d$ the number of genes displaying the top 5% negative post-transcriptional signals and a reduction of at least 2-fold in their exonic (ΔEx) fractions, and that are also putatively targeted by significantly upregulated miRNAs (N = 21 for *AL-T0* vs *AL-T2* and N = 25 for *obese* vs *lean*, **Tables 1** and **2**). Subsequently, the enrichment of the given subset is computed by following three steps: (i) For each row ($i = 1..d$) of $D$, compute $s_i. = \sum_{l=1}^{k} \frac{d_{ik}}{d}$ , i.e. $i \in i_B$. (ii) Taking into account the matrix $M[i_B, .]$, which considers only the rows of $M$ corresponding to the genes of interest and all of its columns ($p$), compute $t_i. = \sum_{l=1}^{p} \frac{m_{il}}{p} \; \forall i \in i_B$. (iii) $s_i$. and $t_i$. are vectors of size $d$, then, the CES value for each gene $i \in i_B$ is computed as the ratio between $s$ and $t$, $CES_i = \frac{s_i}{t_i}$.

Additionally, we defined random sets of genes with the same length as that of genes with top 5% negative post-transcriptional signals and putatively targeted by significantly upregulated miRNAs, i.e. N = 21 for *AL-T0* vs *AL-T2* and N = 25 for *obese* vs *lean* contrasts, and the process described above was repeated iteratively (N = 1,000). We then computed the CES values for all resulting random sets of genes using their exonic and intronic fractions and they were considered as a control set with no expected enrichment in covariation patterns (CES ≈ 1).

### Expression analyses of miRNAs and putative mRNA targets by qPCR

Total RNA extracted from adipocytes of the UNIK minipigs (according to BMI profiles, **Table S1**) had a better quality than RNA isolated from the skeletal muscle samples, so it was subsequently employed for qPCR verification. The same 5 *obese* and 5 *lean* animals used for RNA-seq and small RNA-seq analyses (Jacobsen *et al.* 2019; Sørensen

2014) were employed for qPCR profiling (**Table S1**), with the exception of pig 572, for which no RNA was left. This particular individual was replaced by pig 503, which had the closest BMI profile within the *obese* group of animals (**Table S1**).

Five mRNAs (*LEP*, *OSBPL10*, *PRSS23*, *RNF157* and *SERPINE2*) among those with the top negative 5% post-transcriptional signal (**Table 2**) were selected for qPCR analyses. Accordingly, three of the most significantly upregulated miRNAs were also profiled (ssc-miR-92b-3p, ssc-miR-148a-3p and ssc-miR-214-3p, **Table S10**). Primers were collected from available stocks as reported in previous studies (Mentzel *et al*. 2016; Nygard *et al*. 2007) or designed using the PRIMER3 software within the PRIMER-BLAST tool (Untergasser *et al*. 2012), considering exon–exon junction spanning primers for poly-exonic candidates and ensuring multiple transcript capture whenever possible (**Table S2**).

For mRNA profiling, 400 ng of total RNA were used for cDNA synthesis in duplicate. Briefly, 0.5 μl Improm-II reverse transcriptase (Promega), 0.25 μg 3:1 mixture of random hexamers/OligodT (Sigma-Aldrich), 2 μl 5× ImProm-II buffer, 10 units RNasin Ribonuclease inhibitor (Promega), 2.5 mM $MgCl_2$ and 2mM dNTPs were mixed with 400 ng total RNA in a final reaction volume of 10 μl following manufacturer's instructions. Each reaction was incubated during 5 min at room temperature, 1 h at 42°C and 15 min at 70°C for enzyme inactivation. A negative control (RT) without reverse transcriptase reaction was included. The same RNA samples used for mRNA profiling were subsequently processed for miRNA profiling using 50 ng of total RNA for cDNA synthesis in duplicate and following previous protocols (Balcells *et al*. 2011) with modifications as described in Cirera & Busk (2014). Primers for miRNA qPCR amplification were designed using the miRprimer software (Busk 2014). The synthesized cDNAs were then diluted in a 1:16 proportion for mRNAs and 1:8 for miRNAs and subsequently used for qPCR analyses. Two reference genes, *TBP* and *ACTB* (Nygard *et al*. 2007), plus two highly expressed non-deregulated miRNAs were used for normalization (ssc-let-7a and ssc-miR-23a-3p).

Subsequently, qPCR for mRNA and miRNA genes was performed on a MX3005P machine (Stratagene, USA) using the QuantiFast Multiplex PCR protocol kit (Qiagen, Germany). Briefly, diluted cDNAs were mixed with 5 µl of 2× QuantiFast Multiplex master mix (Qiagen, Germany) and 10 µM of each primer (**Table S2**) were mixed in a final volume of 10 µl. Cycling conditions were: 95 °C for 5 min followed by 40 cycles of 95 °C for 10 sec and 60 °C for 30 sec. Melting curve analyses (60 °C to 99 °C) were performed after completing amplification reactions to ensure the specificity of the assays. Data were processed with the MxPro qPCR associated software. Assays were considered successful when: (i) the melting curve was specific (1 single peak) and (ii) the samples had Cq values < 33 cycles (i.e. sufficiently expressed to be considered biologically functional).

Efficiency of each primer assay was estimated from the log–linear fraction of their corresponding standard curves. Amplification efficiency ranging between 80–110% was considered acceptable with $R^2 > 0.98$. Pre-processing of raw quantification cycle (Cq) values was performed with the GenEx v.6 software (MultiD, Analyses AB). Briefly, raw expression data for each successfully amplified gene were corrected by taking into account PCR efficiency, and Cq values were normalized using mRNA and miRNA reference genes previously evaluated with the *geNorm* algorithm (Vandesompele *et al*. 2002). Duplicates for each sample were subsequently averaged and relative quantities were scaled to the lowest expressed sample in each assay, and $\log_2$ transformed before statistical analyses. The significance of differential expression between *obese* and *lean* animals was estimated with the limma empirical Bayes procedure using the *ebayes* function of the *limma* R package (Ritchie *et al*. 2015) incorporating a mean-variance trend modeling the relationship between variance and gene signal intensity and correcting for sex (**Table S1**) as fixed covariate.
